# Differential Signaling Mediated by ApoE2, ApoE3, and ApoE4 in Human Neurons Parallels Alzheimer’s Disease Risk

**DOI:** 10.1101/460899

**Authors:** Yu-Wen Alvin Huang, Bo Zhou, Amber M. Nabet, Marius Wernig, Thomas C. Südhof

## Abstract

Apolipoprotein E (ApoE) mediates clearance of circulating lipoproteins from blood by binding to ApoE receptors. Humans express three genetic variants, ApoE2, ApoE3, and ApoE4, that exhibit distinct ApoE receptor binding properties. In brain, ApoE is abundantly produced by activated astrocytes and microglia, and three variants differentially affect Alzheimer’s disease (AD), such that ApoE2 protects against, and ApoE4 predisposes to the disease. A role for ApoE4 in driving microglial dysregulation and impeding Aβ clearance in AD is well documented, but the direct effects of three variants on neurons are poorly understood. Extending previous studies, we here demonstrate that ApoE variants differentially activate multiple neuronal signaling pathways and regulate synaptogenesis. Specifically, using human neurons cultured in the absence of glia to exclude indirect glial mechanisms, we show that ApoE broadly stimulates signal transduction cascades which among others enhance synapse formation with an ApoE4>ApoE3>ApoE2 potency rank order, paralleling the relative risk for AD conferred by these variants. Unlike the previously described induction of *APP* transcription, however, ApoE-induced synaptogenesis involves CREB rather than cFos activation. We thus propose that in brain, ApoE acts as a glia-secreted paracrine signal and activates neuronal signaling pathways in what may represent a protective response, with the differential potency of ApoE variants causing distinct levels of chronic signaling that may contribute to AD pathogenesis.

## Introduction

ApoE is a major apolipoprotein of circulating lipoproteins that is mainly produced by hepatocytes and mediates binding of lipoproteins to LDL-and VLDL-receptors, which are the major peripheral ApoE receptors (1, 2). Humans but not other animals express three allelic ApoE variants, ApoE2, ApoE3, and ApoE4, that differ at two amino acid residues and exhibit distinct ApoE receptor binding properties. Specifically, ApoE2 binds much less strongly to ApoE receptors than the other two variants, whereas at least under some conditions ApoE4 appears to bind ApoE receptors more effectively than ApoE3 (3–5). Likely due to these differences in receptor binding properties, homozygosity for ApoE2 predisposes to familial dysbetalipoproteinemia, while homozygosity for ApoE4 causes hypercholesterolemia and atherosclerosis (6, 7). In addition to hepatocytes in liver, activated astrocytes and microglia in brain also abundantly produce ApoE (8, 9). Strikingly, carriers of an ApoE4 allele exhibit a greatly increased risk for Alzheimer’s disease (AD) even as heterozygotes, such that ApoE4 is the most important genetic risk factor for AD (10). Carriers of the ApoE2 allele, in contrast, display a decreased risk for AD (11).

Although its connection to AD suggest that ApoE has a significant role in brain, the function of ApoE in brain and the mechanistic relation of ApoE4 to AD is incompletely understood. Human patients who lack ApoE expression because of mutations in both of their ApoE alleles suffer from severe dysbetalipoproteinemia, but appear to be cognitively normal and do not exhibit major neurological impairments (12–14). This observation suggests that ApoE in brain has no constitutive essential function, and that it is not required to prevent AD or other neurodegenerative processes. However, ApoE could serve as a protective signal in brain that only assumes functional relevance when a repair reaction or protective response is elicited, a possibility that is supported by the finding that activation of microglia and astrocytes induces ApoE expression (15, 16). Informed by these data, three major hypotheses that are not mutually exclusive have been advanced to account for the fact that ApoE2 protects against AD pathogenesis, whereas ApoE4 promotes AD pathogenesis.

First, it has been shown that ApoE is important for the clearance of amyloid-β (Aβ) peptides, which in turn are thought to be central players in AD pathogenesis (17–20). This observation suggests that ApoE4 promotes AD pathogenesis by increasing the concentration of pathogenic Aβ, but the precise mechanisms involved remain uncertain, as does the actual role of Aβ in AD pathogenesis.

Second, recent studies revealed that another major risk factor for AD is a genetic variation in TREM2, a microglial surface receptor (21–23), and that ApoE binds to TREM2 (24–26). ApoE binding to TREM2 activates an intracellular signal transduction cascade consistent with a regulatory function in microglia (27), and ApoE4 specifically causes amyloid-dependent microglial dysfunction (28, 29). Thus, ApoE4 could promote AD pathogenesis by impeding normal microglial function. However, TREM2 lacks the classical ApoE-binding sequence motif that was first defined in the LDL-receptor (30), raising the question of how exactly ApoE binds to TREM2. Moreover, microglia themselves produce copious amounts of ApoE (31–33), suggesting that ApoE might serve as an autocrine signal in microglia, which in turn prompts the question of the biological utility of such a mechanism.

Third, ApoE2, ApoE3, and ApoE4 may bind to ApoE receptors on neurons and differentially activate an intracellular response, the pathogenic consequences of which might arise only after long-term chronic activation. This hypothesis is based on the potent activation of intracellular signaling by ApoE receptors in neurons (34–36) and the induction of APP and Aβ synthesis by ApoE with a potency rank order of ApoE4>ApoE3>ApoE2 (37). However, this effect of ApoE on neurons is only observed in neurons in the absence of glia and serum because astrocytes themselves produce copious amounts of ApoE and serum naturally contains high concentrations of ApoE (37). The dependence of this activation of neuronal signaling by ApoE on the precise experimental conditions may explain questions raised regarding the reproducibility of this effect (38).

All three hypotheses regarding the mechanism of ApoE4’s role in AD pathogenesis are intriguing, but lack sufficient experimental support to be considered definitive. In the present study, we focused on the third hypothesis because it addresses a specifically neuronal function. If ApoE indeed mediated a protective response that is not required in an unchallenged brain, activation of neuronal signaling by ApoE could be part of an inducible repair response, but chronic hyperactivation of such a response might be deleterious. Indeed, we here show that ApoE receptor activation broadly stimulates multiple signal transduction cascades in neurons that govern many cellular responses, including the enhancement of synapse formation without dendritic arborization changes. ApoE enhanced synapse formation with an ApoE4>ApoE3>ApoE2 potency rank order in a manner requiring MAP kinase activation. In contrast to the previously described induction of APP transcription by ApoE (37) but consistent with an earlier study (35), ApoE-induced synapse formation involved activation of CREB. Thus, in addition to probable autocrine effects of microglial ApoE in brain, we propose that ApoE acts as a paracrine signal to broadly activate neuronal signaling pathways in what may represent a protective biological response.

## Results

### ApoE binding to neuronal receptors stimulates an array of signaling pathways

A possible direct activation of neuronal signaling by ApoE has been reported by us and others (34–37), but could not always be reproduced (38). Given growing concerns about reproducibility, validation of scientific results is arguably more important than the prominent question of whether a particular result is physiologically relevant. To address this issue directly for the activation of neuronal signaling by ApoE, we embarked on a replication effort using a rigorous experimental design. We first validated that ApoE is not produced at significant levels by human neurons differentiated from ES cells, even though ES cells express surprisingly high levels of ApoE (Supplemental Figure 1, A and B). Moreover, we analyzed recent mouse brain single-cell RNAseq data (39, 40) and confirmed that in mice ApoE is also expressed at much lower levels in neurons than in astrocytes and microglia (Supplemental Figure 1, C and D). These results argue against the notion that ApoE functions as a major neuronal protein that is pathogenic when misfolded in the cytoplasm of neurons (38), but are consistent with the idea that ApoE is produced as a signaling factor by glia.

We next tested in a ‘double blind’ fashion the signaling effects of ApoE on human neurons that were cultured in the absence of serum on mouse embryonic fibroblasts (MEFs, which produce ApoE only at very low levels; Supplemental Figure 1, A and B). Two scientists (Y.A.H. and A.M.N.) each independently produced ApoE2, ApoE3, and ApoE4 proteins in transfected HEK293 cells; these protein preparations were then randomly assigned numbers to ‘anonymize’ them by a third scientist, and the original two scientists used the anonymized samples to test whether the effect of ApoE variants on ERK phosphorylation and APP levels were reproducible. ApoE2, ApoE3, and ApoE4 proteins synthesized in HEK293 cells are secreted into the supernatant without significant differences in abundance and are pelleted by ultracentrifugation by similar g forces, suggesting that recombinant ApoE2, ApoE3, and ApoE4 produced in transfected HEK293 cells are biochemically similar (Supplemental Figure 2,A and B). Double-blinded application of the two separate ApoE preparations by the two experimenters to human neurons yielded essentially the same results as those described earlier (37), namely a stimulation of ERK1/2 phosphorylation and an increase in the levels of APP and DLK proteins, all of which were induced with a potency rank order of ApoE4>ApoE3>ApoE2 (Figure 1, A and B; Supplemental Figure 3A). The effect size differed between experimenters but not between ApoE preparations, probably because of person-to-person variabilities in the technically complex preparation of human neurons cultured on MEFs without serum supplementation. Nevertheless, the effects were significant for both ApoE preparations for both experimenters.

**Figure 1.**
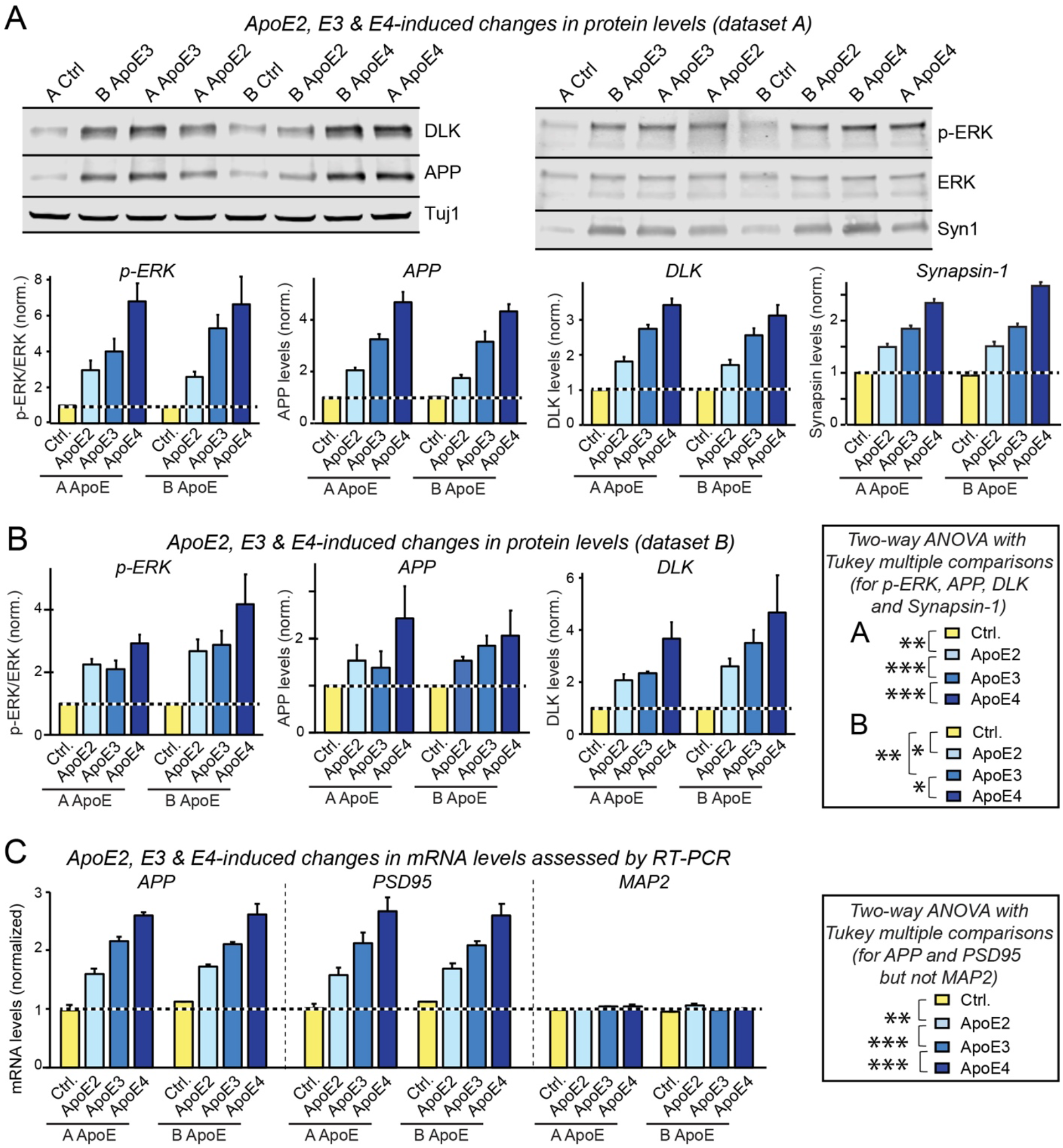
ApoE-induced stimulation of ERK phosphorylation and induction of DLK and APP proteins is reproduced by multiple experimenters using anonymized independent ApoE preparations. Data are from human neurons cultured on MEFs in the absence of serum; recombinant ApoE (10 μg/mL, produced in HEK293 cells, see Figure S1F) or control solutions were added at day 10 (D10), and neurons were analyzed at D12 as indicated. ApoE and control solutions were prepared by two experimenters (‘A ApoE’ and ‘B ApoE’), anonymized, and used by the same experimenters (‘dataset A’ and ‘dataset B’). (**A**) Dataset A showing that ApoE induces an increase in ERK phosphorylation and in the levels of DLK, APP, and synapsin-1 (Syn1) proteins with an ApoE4>ApoE3>ApoE2 potency rank order (top, representative immunoblots; bottom, summary graphs). Protein levels measured using fluorescent secondary antibodies were normalized for the Tuj 1 signal examined on the same blots as an internal standard, and additionally for the levels observed in control neurons. Because ApoE solutions were anonymized, samples on the immunoblot are not in a logical order. (**B**) Dataset B showing that ApoE induces an increase in ERK phosphorylation and in the levels of DLK and APP proteins with an ApoE4>ApoE3>ApoE2 potency rank order (synapsin-1 was not analyzed). Only summary graphs are shown; for representative immunoblots, see Figure S2. (**C**) ApoE2, ApoE3, and ApoE4 increase neuronal APP and PSD95 but not MAP2 gene expression as assessed by mRNA measurements in human neurons treated with the two independently produced ApoE preparations. mRNA levels were normalized for those of MAP2 as an internal standard that is specific for neurons to avoid monitoring indirect effects of neuronal differentiation which in itself results in an increase in MAP2 mRNA levels, and for the levels observed in the absence of ApoE (=1.0). Data in bar graphs are means ± SEM (n ≥ 3 independent experiments); statistical significance was evaluated by two-way ANOVA with Tukey multiple comparisons (*, p<0.05, **, p<0.01; ***, p<0.001) as detailed in the boxes. Note that in all analyses in which ApoE has an effect on a measured parameter, ApoE3 is always significantly more potent than ApoE2 and less potent than ApoE4.

In some of the replication experiments, we included analysis of synapsin-1 protein as a synaptic marker of human neurons. We were surprised to observe an increase in synapsin-1 levels upon ApoE treatment (Figure 1A). To confirm this finding, we measured mRNA levels of APP and of another synaptic marker, PSD95, in a parallel experiment, using the pan-neuronal marker MAP2 as a control. Again, we observed a significant induction of PSD95 levels by ApoE (Figure 1C). These results suggested the possibility that ApoE may produce a broader signal transduction response than we had originally envisioned (37). To test this possibility, we surveyed in the human neurons the effect of ApoE on the phosphorylation of four key signal transduction proteins, ERK1/2, Akt, Src, and JNK (Figure 2A). In these experiments, we analyzed both neurons cultured on MEFs and neurons cultured on matrigel alone in the absence of any cellular support in order to ensure that the observed effects did not depend on the presence of the MEFs. We found ApoE strongly stimulated phosphorylation of Akt and Src in human neurons in addition to that of ERK, again with a potency rank order of ApoE4>ApoE3>ApoE2, but that phosphorylation of JNK was not affected. All of these effects were observed similarly with or without MEFs as a cellular substrate for the neuronal culture (Figure 2A), and were abolished by the ApoE receptor blocking protein RAP, demonstrating that they were induced by ApoE receptor binding (Figure 2B).

**Figure 2.**
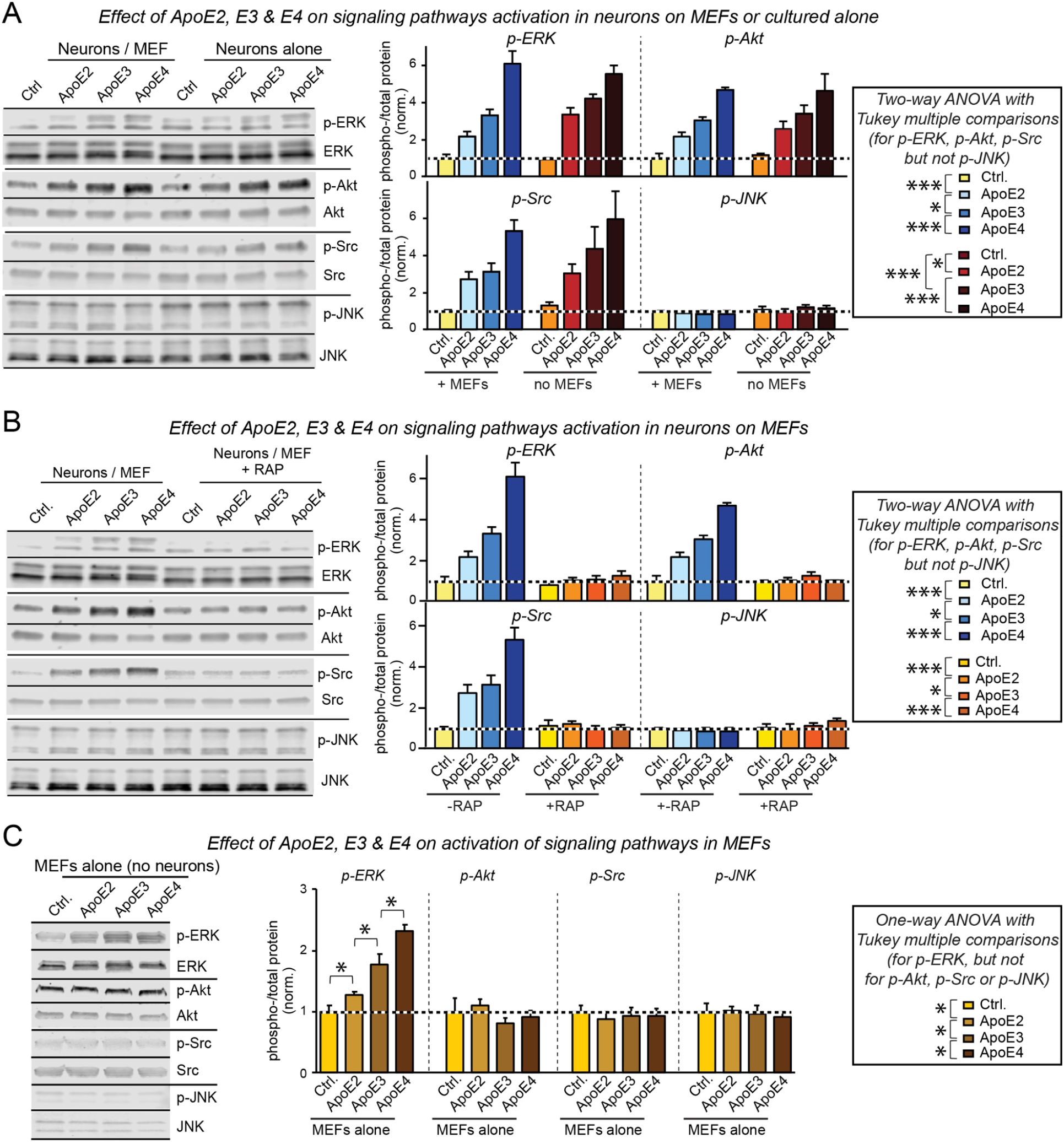
ApoE potently activates multiple signaling pathways in human neurons with an ApoE4>ApoE3>ApoE2 potency rank order, but stimulates only ERK phosphorylation in MEFs. Data are from human neurons cultured on MEFs or matrigel or from only MEFs cultured in the absence of serum; recombinant ApoE (10 μg/ml, produced in HEK293 cells, see Supplemental Figure 1F) or control solutions were added for 1 h at D10 before the indicated phosphorylation events were analyzed by immunoblotting. (**A**) ApoE induces phosphorylation of ERK, Akt and Src but not of JNK in human neurons cultured on MEFs or matrigel with an ApoE4>ApoE3>ApoE2 potency rank order (left, representative immunoblots; right, summary graphs). (**B**) ApoE3-induced ERK-, Akt-and Src-phosphorylation are prevented by the ApoE receptor blocking protein RAP (50 μg/ml, applied 30 min before the 1 h ApoE incubation at D10; left, representative immunoblots; right, summary graphs). (**C**) ApoE induces phosphorylation of ERK, but not Akt, Src or JNK in MEFs that were cultured in the absence of human neurons and treated with ApoE as in A (left, representative immunoblots; right, summary graphs). Data are means ± SEM (n>3 independent experiments); statistical significance (*, p<0.05; ***, p<0.001) was evaluated with two-way (**A, B**) or one-way ANOVA (**C**) and selected Tukey’s post-hoc multiple comparisons as indicated.

Is the signal transduction response to ApoE observed in human neurons a general cellular response? To examine this question, we tested ERK, Akt, Src, and JNK phosphorylation as a function of ApoE in MEFs cultured without neurons. Indeed, ApoE also induced ERK phosphorylation in MEFs, again with a rank-potency order of ApoE4>ApoE3>ApoE2, but had no effect on Akt or Src phosphorylation (Figure 2C). Thus, ApoE may generally stimulate signal transduction in cells consistent with previous studies (34–36), with cell-type specific differences in the response patterns.

### ApoE activates synapse formation

The unexpected ApoE-induced increase in synaptic markers (Figure 1) is reminiscent of the observation that ApoE-containing lipoprotein particles stimulate synapse formation (41), prompting us to examine whether synapse formation in human neurons cultured on MEFs is stimulated by ApoE. Compared to neurons cultured on glia, neurons cultured on MEFs exhibit a ~3-fold decrease in synapse density without a change in neuronal soma size or dendritic arborization, presumably because glia secrete potent synaptogenic factors (Figure 3, A to F). Addition of ApoE to neurons cultured on MEFs in the absence of serum induced a 1.5-to 2-fold increase in synapse density, with a rank-potency order of ApoE4>ApoE3>ApoE2 (Figure 3, A to F; Supplemental Figure 4, A and B). This increase was abolished in the presence of the ApoE receptor blocker RAP, whereas RAP had no significant effect on synapse density in neurons cultured on glia, which secrete other synaptogenic factors besides ApoE (Figure 3, A to F). In these experiments, we analyzed synapse density by staining the neurons for the presynaptic marker protein synapsin-1, but we observed a similar effect when we analyzed synapse density using Homer1 as a postsynaptic marker (Figure 3, G to J). Moreover, when we performed similar experiments at an older age of cultured neurons (DIV23-25) we observed a comparable effect (Supplemental Figure 4, A and B).

**Figure 3.**
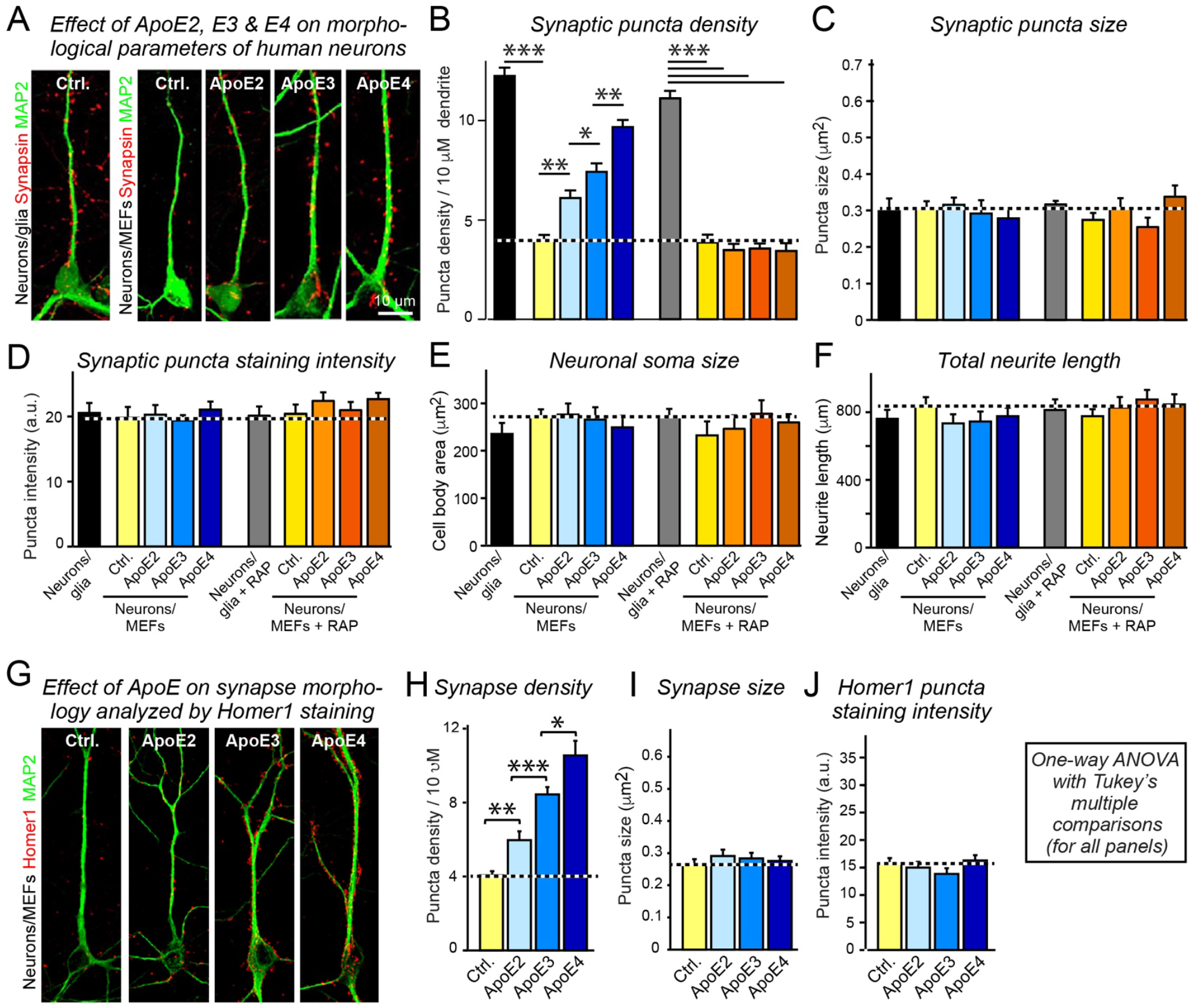
ApoE increases synapse formation in human neurons cultured on MEFs in a manner inhibited by the ApoE-receptor blocking protein RAP, with a rank potency order of ApoE4>ApoE3>ApoE2. Data were from human neurons cultured on MEFs in the absence of serum; recombinant ApoE (10 μg/ml, produced in HEK293 cells) or control solutions were added at D10, and neurons were analyzed at D16. (**A**) Representative images of human neurons analyzed by double-immunofluorescence labeling for MAP2 (a dendritic marker) and synapsin (a presynaptic marker). (**B–F**) Summary graphs of the synapse density (**B**), synaptic puncta size (**C**), synapsin puncta staining intensity (**D**), cell body area (E), and total neurite length (**F**) in human neurons cultured in the absence and presence of ApoE receptor blocking protein RAP (50 μg/ml). (**G**) Representative images of human neurons labelled by dual immunofluorescence staining for MAP2 and Homer1 (a post-synaptic marker). (**H**–**J**) Summary graphs of the synaptic Homer1 puncta density (**H**), Homer1 puncta size (**I**), and Homer1 puncta staining intensity (**J**). Data are means ± SEM (n≥3 independent experiments); statistical significance (*, p<0.05, **, p<0.01; ***, p<0.001) was evaluated with one-way ANOVA with Tukey’s multiple comparisons.

The measurements of synapsin-1 protein and PSD95 mRNA that we performed in the context of the replication experiments (Figure 1) suggested that ApoE may stimulate synaptic gene expression in promoting synapse formation. Consistent with this hypothesis, we found that ApoE, again with a potency rank order of ApoE4>ApoE3>ApoE2, enhanced expression of all synaptic genes tested both at the mRNA and the protein levels (Figure 4, A and B). As for all effects of ApoE on human neurons, this enhancement was abolished by the ApoE receptor blocker RAP, and was dose-and time-dependent (Supplemental Figure 4C).

Previous experiments suggested that ApoE-containing lipoprotein particles stimulate synapse formation by delivering cholesterol (41). To test this hypothesis, we examined whether ApoE made in bacteria that are incapable of synthesizing cholesterol would also stimulate synapse formation. ApoE was produced efficiently in bacteria (Supplemental Figure 5A). Importantly, bacterial ApoE was as efficacious as HEK293 cell-produced ApoE in stimulating synaptic gene expression (Figure 4C), ruling out cholesterol as a major agent. Moreover, we examined the possibility that the action of ApoE on synaptic gene expression was specific for human neurons produced from H1 ES cells, but detected a similar effect in neurons produced from two different lines of induced pluripotent stem (iPS) cells (Supplemental Figure 6, A and B). Finally, since ApoE stimulates the MAP kinase pathway in MEFs similar to neurons (Figure 2C), we asked whether ApoE-induced synapse formation in neurons cultured on MEFs could be an indirect effect of the activation of MEFs. However, ApoE was as effective at stimulating synaptic gene expression in neurons cultured on an inanimate matrigel support as neurons cultured on MEFs in, indicating that ApoE acts directly on the neurons (Figure 4D). In all of these experiments, we observed the same differential efficacy with a potency rank order of ApoE4>ApoE3>ApoE2.

**Figure 4.**
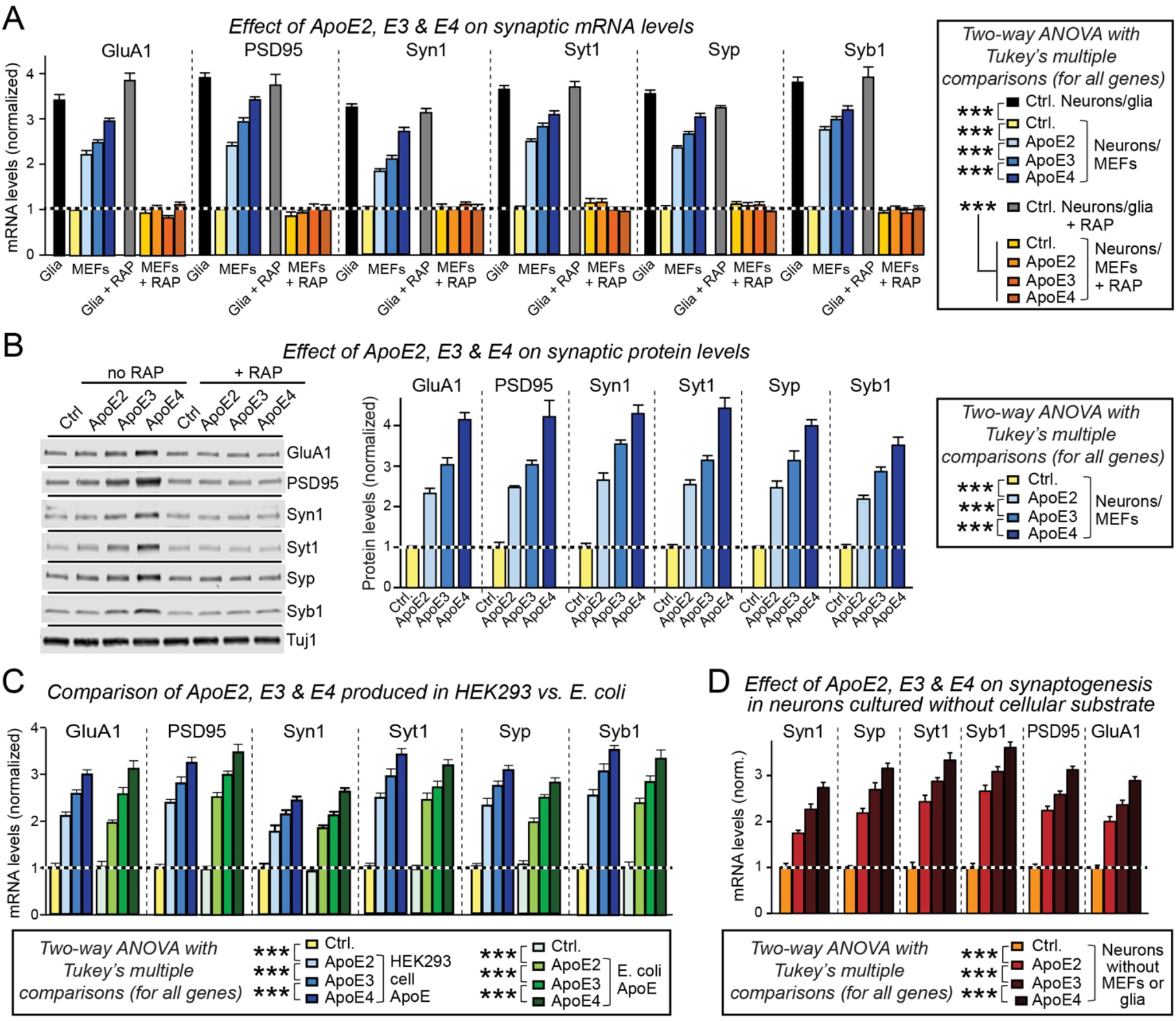
ApoE stimulates synaptic gene transcription independent of its lipidation state or culture substrate, with a rank potency order of ApoE4>ApoE3>ApoE2. Data are from human neurons cultured on MEFs in the absence of serum; recombinant ApoE (10 μg/ml, produced in HEK293 cells or bacteria as indicated) or control solutions were added at D10 with or without RAP (50 μg/ml), and neurons were analyzed at D12. (**A**) ApoE increases synaptic protein mRNA levels with an ApoE4>ApoE3>ApoE2 potency rank order; blocking ApoE receptors with RAP abolishes ApoE-induced synaptic mRNA increases. mRNA levels were measured by human-specific qRT-PCR with MAP2 as internal standard to correct for the progressive neuronal maturation during the experiment, and plotted normalized to controls (abbreviations: GluA1, glutamate receptor subunit-1; Syt1, synaptotagmin-1; Syp, synaptophysin; Syn, synapsin; Syb1, synaptobrevin-1). (**B**) ApoE increases synaptic protein levels with an ApoE4>ApoE3>ApoE2 potency rank order (left, representative immunoblots; right, summary graphs of protein levels normalized for Tuj1 as an internal standard). (**C**) Recombinant ApoE produced in bacteria (E. coli) and HEK293 cells are equally potent in increasing synaptic protein mRNA levels in neurons cultured in the absence of glia or serum. Analyses were performed as in A. (**D**) ApoE increases synaptic protein mRNA levels with an ApoE4>ApoE3>ApoE2 potency rank order also when human neurons are cultured on matrigel without cellular support. Summary graphs show the indicated synaptic protein mRNA levels measured by human-specific quantitative RT-PCR with MAP2 as internal standard, and normalized to the untreated control. mRNA levels are normalized to the control and to MAP2 as an internal standard. Data are means ± SEM (n≥3 independent experiments); statistical significance (***, p<0.001) was evaluated by two-way ANOVA with Tukey’s multiple comparisons.

The significant increase in synapse numbers induced by ApoE should effectuate an increase in synaptic transmission. We investigated this conjecture by recording from neurons that were cultured on MEFs in the absence of glia or serum and were incubated in either control medium or medium containing ApoE2, ApoE3, or ApoE4. None of the ApoE variants had an effect on the capacitance or input resistance of the neurons (Figure 5, A and B), but all variants augmented the amplitude of evoked EPSCs 1.5-to 3-fold, again with a potency rank order of ApoE4>ApoE3>ApoE2 (Figure 5, C and D). Furthermore, ApoE similarly increased the frequency of spontaneous mEPSCs recorded in the presence of tetrodotoxin without a significant effect on mEPSC amplitudes (Figure 5, E to G). Thus, ApoE stimulates formation of functional synapses in neurons cultured in the absence of glia or serum.

**Figure 5.**
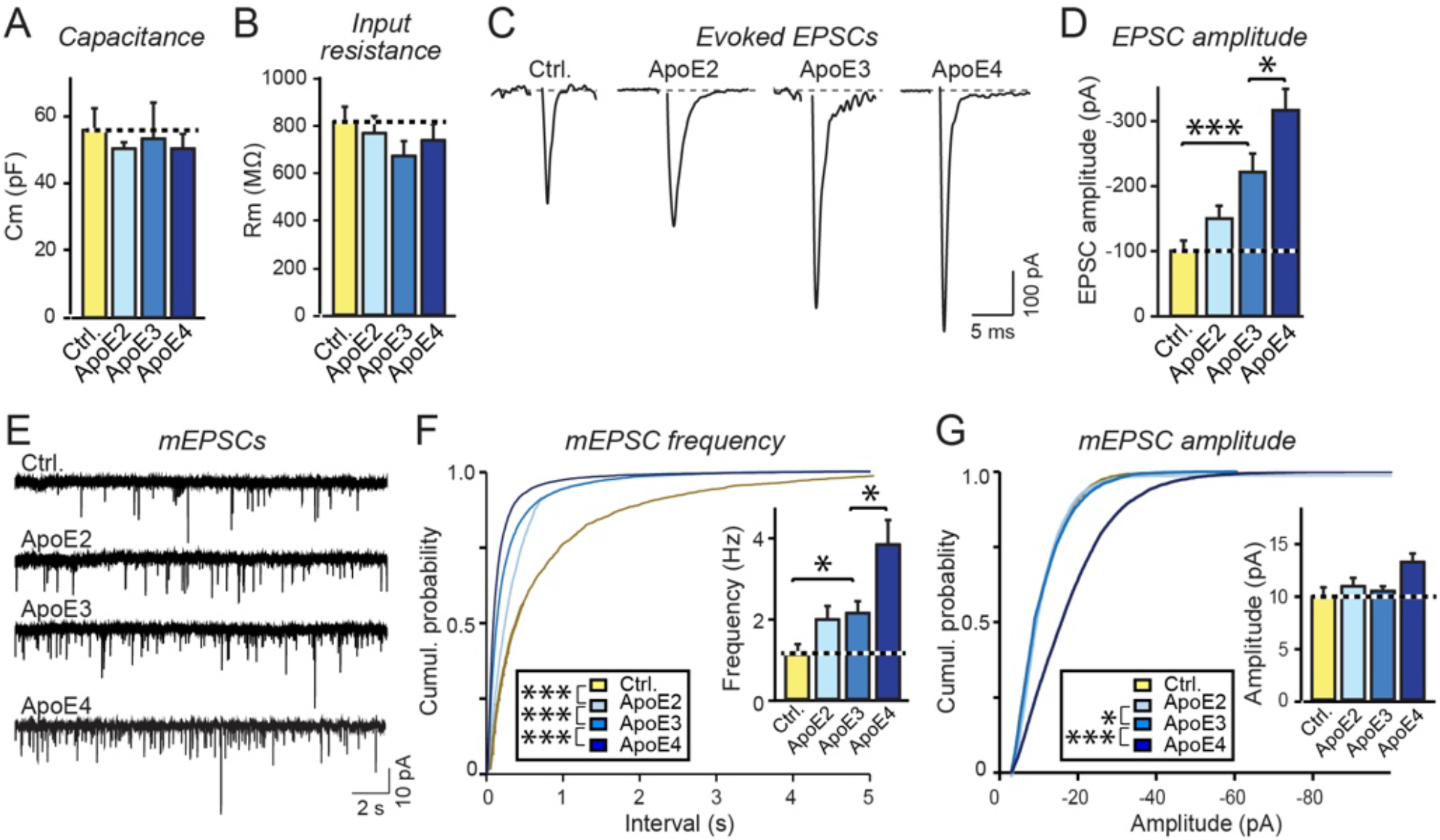
ApoE increases excitatory synaptic strength but does not change the intrinsic electrical properties of human neurons, with ApoE4 being more and ApoE2 being less efficacious than ApoE3. Data are from human neurons cultured on MEFs in the absence of serum; recombinant ApoE (10 μg/ml) or control solutions were added at D10, and neurons were analyzed at D23-25. (**A & B**) Summary graphs of membrane capacitance (**A**) and resistance (**B**). (**C**) Representative traces of evoked EPSCs. (**D**) Quantification of evoked EPSC amplitudes in human neuronal cultures. (**E**) Representative traces of mEPSCs from human neurons. (**F** & **G**) Summary plots and graphs of the frequency (**F**) and amplitudes (**G**) of mEPSCs. Quantifications are shown both as cumulative probability plots and as bar diagrams (inserts). Electrophysiological recordings were done in parallel with morphological analyses (Supplemental Figure 4, B and C). Statistical significance (*p < 0.05; ***p<0.001) was performed with one-way ANOVA with post-hoc Tukey multiple comparisons for all bar diagrams, presented as means ± SEM (n = 31–38 in 3 batches for evoked EPSCs; n = 32–39 in 5 batches for mEPSCs); Kolmogorov-Smirnov Test was performed for cumulative probability plots in (**F**) and (**G**).

### ApoE-induced synapse formation is mediated by ERK activation

We previously showed that ApoE enhances APP transcription by activating MAP kinases, but our present data suggest that ApoE additionally stimulates multiple other signaling pathways (Figure 2). We thus tested whether ApoE induced synaptic gene expression is sensitive to a general MAP kinase inhibitor, U0126, or to a PI3-kinase inhibitor, Wortmannin. U0126 blocked the ApoE-induced increase in synaptic gene expression more efficiently than the ApoE receptor blocker RAP (Figure 6A), whereas Wortmannin had no effect (Figure 6B). Moreover, ApoE acted via the same MAP kinases in stimulating synapse formation as in enhancing APP transcription because shRNA targeting DLK, a MAP-kinase kinase kinase that is essential for the ApoE-induced increase in APP (37), also abolished the increase in synaptic gene expression, whereas overexpression of DLK constitutively enhanced synaptic gene expression (Figure 6, A and B). Consistently, overexpression of MBIP, an inhibitor of DLK (42), abolished the effect of ApoE and even decreased the baseline expression of synaptic genes (Figure 7, A and B). Furthermore, CRISPR/Cas9 directed to MKK7, a MAP-kinase kinase downstream of DLK, also abolished the effect of ApoE and lowered baseline synaptic gene expression (Figure 7, A and B); again, overexpression of MKK7 overrode the effect of MKK7 CRISPR disruption and constitutively enhanced synaptic gene expression similar to overexpression of DLK. Thus, the same MAP kinase pathway is required for both ApoE-induced synapse formation and ApoE-induced enhancement of *APP* transcription.

**Figure 6.**
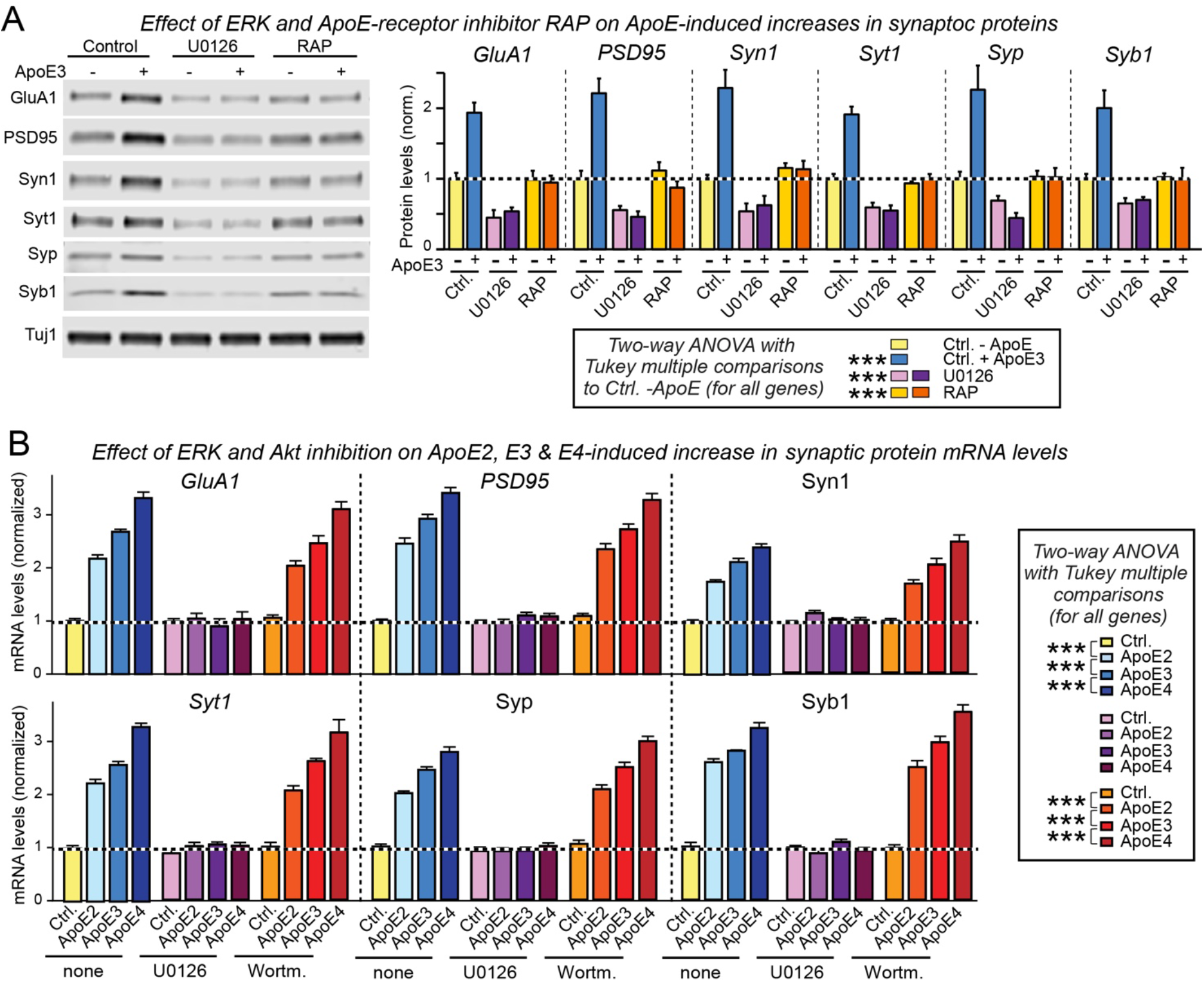
ApoE induction of synaptic gene expression is blocked by the MAP-kinase inhibitor U0126 and the ApoE-receptor blocking protein RAP. Data are from human neurons cultured on MEFs in the absence of serum; recombinant ApoE (10 μg/ml) or control solutions without or with U0126 (50 μM), RAP (50 μg/ml), or Wortmannin (0.1 μM) were added at D10, and neurons were analyzed at D12. (**A**) ApoE3-induced increases in synaptic protein levels is abolished by the MAP-kinase inhibitor U0126 and the ApoE receptor blocking protein RAP (left, representative immunoblots; right, summary graphs of the indicated synaptic protein levels). (**B**) ApoE-induced increases in the levels of mRNAs encoding synaptic proteins is abolished by the MAP-kinase inhibitor U0126 but not by PI-3 kinase inhibitory Wortmannin. mRNA levels are normalized to the control and to MAP2 as an internal standard. Note that the rank potency order of ApoE4>ApoE3>ApoE2 is maintained under all conditions under which ApoE is active. Data are means ± SEM (n≥3 independent experiments); statistical significance (***, p<0.001) was evaluated with two-way ANOVA with Tukey’s multiple comparisons.

**Figure 7.**
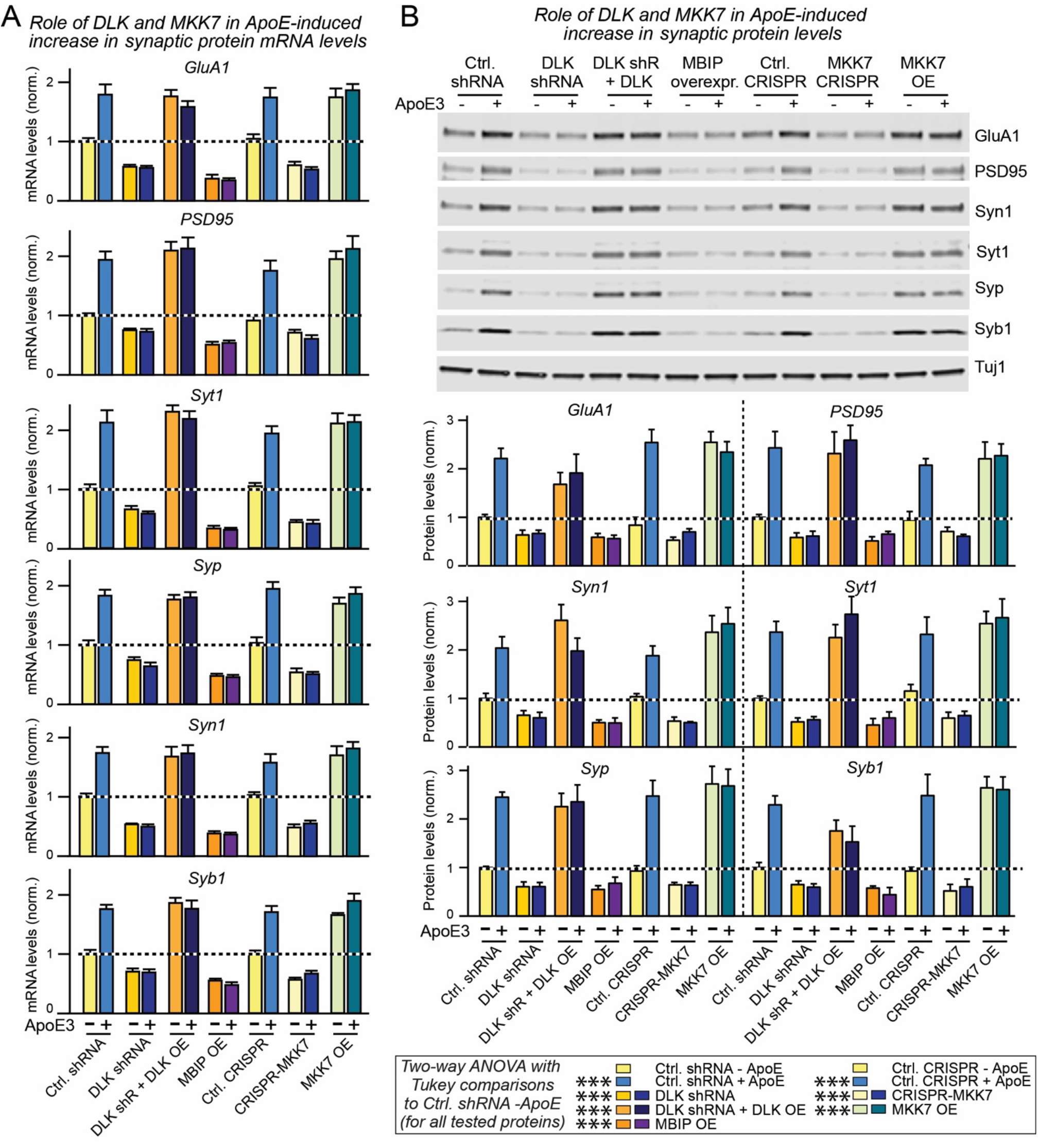
ApoE induction of synaptic gene expression requires the MAP-kinase kinase kinase DLK and the MAP kinase kinase MKK7. Data are from human neurons cultured on MEFs in the absence of serum; the indicated molecular manipulations (DLK knockdowns with shRNAs, CRISPR-suppression of MKK7, and/or overexpression of DLK, MMK7 or MBIP) were initiated by lentiviral transduction at D4, recombinant ApoE3 (10 μg/ml) was added at D10, and neurons were analyzed at D12. (**A**) shRNA-mediated knockdown of DLK, CRISPR-mediated suppression of MKK7 expression, or MBIP-mediated blockage of DLK activity abolish the ApoE-induced increase in synaptic protein mRNA levels, whereas DLK or MKK7 overexpression constitutively increase synaptic protein mRNA levels and render them insensitive to ApoE3 (abbreviations: OE, overexpression; shR, shRNAs). mRNA levels are normalized to the control and to MAP2 as an internal standard. (**B**) shRNA-mediated knockdown of DLK, CRISPR-mediated suppression of MKK7 expression, or MBIP-mediated blockage of DLK activity abolish the ApoE-induced increase in synaptic protein mRNA levels, whereas DLK or MKK7 overexpression constitutively increase synaptic protein mRNA levels and render them insensitive to ApoE3 (top, representative immunoblots; bottom, summary graphs of protein levels normalized to Tuj 1 as an internal standard; abbreviations: OE, overexpression; shR, shRNAs). Data are means ± SEM (n>3 independent experiments); statistical significance (***, p<0.001) was evaluated with two-way ANOVA with Tukey’s multiple comparisons.

### ApoE-induced synapse formation involves CREB but not cFos

ApoE-induced MAP kinase activation stimulates cFos phosphorylation, which in turn enhances APP transcription (37). To explore whether the same pathway operates for ApoE-induced synaptic gene expression that is also mediated by MAP-kinase activation, we examined the effect of a dominant-negative cFos mutant (Figure 8A). Although dominant-negative cFos blocked the effect of ApoE on APP transcription, it had no effect on the ApoE-induced increase in synaptic gene expression (Figure 8A). We therefore tested dominant-negative mutants of two other transcription factors known to be important for neuronal function, CREB and MEF2A. The mutants of human proteins were designed based on validated rodent cDNA sequences (43, 44). Whereas dominant-negative MEF2A had no significant effect on the ApoE stimulation of synaptic gene expression, dominant-negative CREB not only blocked ApoE stimulation of synaptic gene expression, but also decreased baseline levels of synaptic gene expression (Figure 8A). Dominant-negative CREB did not interfere with the ApoE stimulation of APP expression, but appeared to also decrease baseline transcription of *APP* as well.

**Figure 8.**
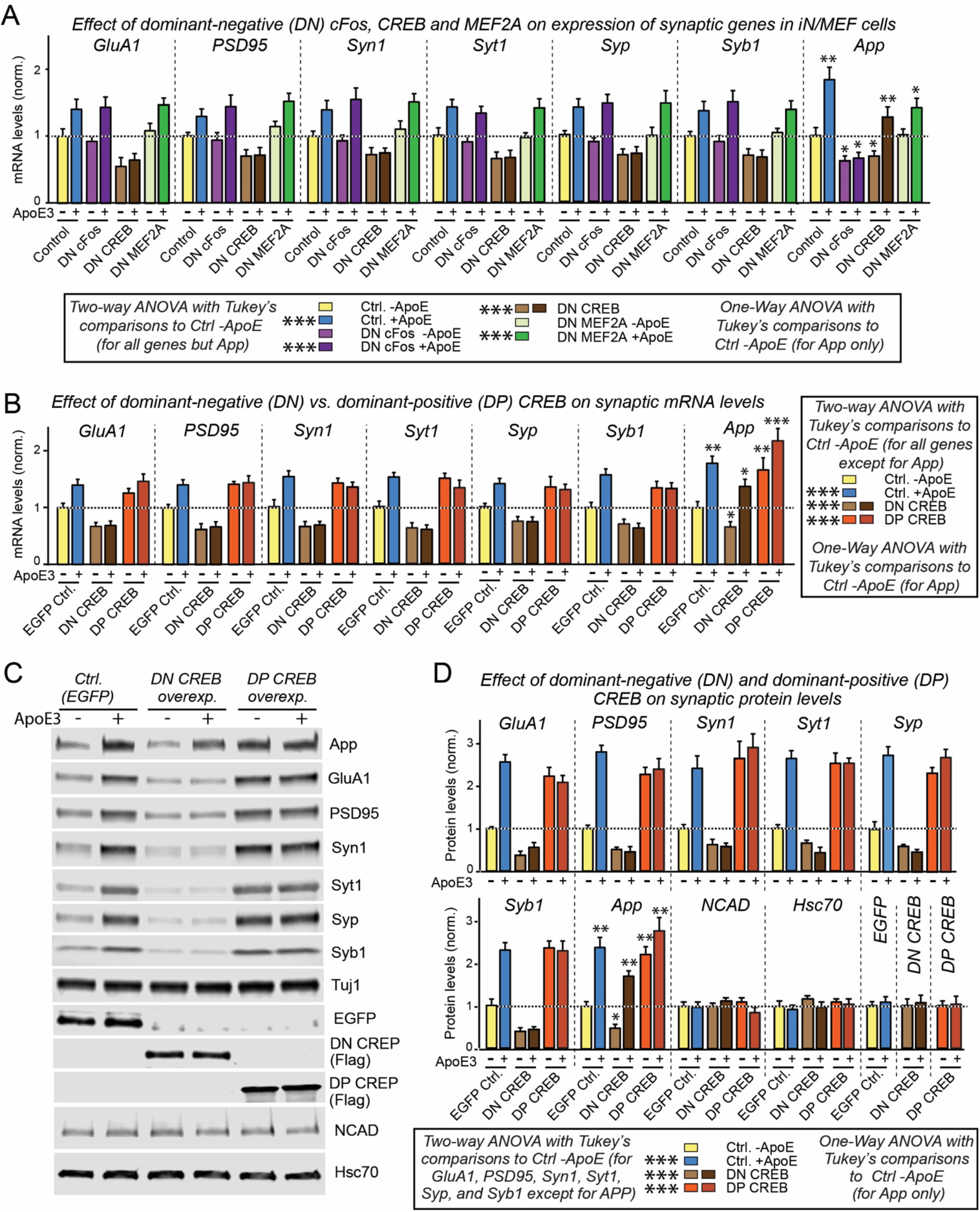
ApoE3-induced increases in synaptic protein expression require CREB but not cFos. Data are from human neurons cultured on MEFs in the absence of serum; the indicated molecular manipulations (overexpression of dominant-negative [DN] mutants of CREB, cFos, or MEF2A, or of a dominant-positive [DP] mutant of CREB) were initiated by lentiviral transduction at D4, recombinant ApoE3 (10 μg/ml) was added at D10, and neurons were analyzed at D12. (**A**) DN-CREB but not DN-cFos or DN-MEF2A inhibits the ApoE3-induced increase in mRNA levels for synaptic proteins, whereas DN-cFOS but not DN-CREB or DN-MEF2A blocks the ApoE3-induced increase in APP mRNA levels. mRNA levels were quantified by qRT-PCR and are shown normalized for those of MAP2 and of controls. (**B**) DN-CREB suppresses baseline levels of synaptic protein and APP mRNAs and abolishes ApoE3-induced increases of synaptic protein but not APP mRNAs; DP-CREB, in contrast, constitutively increases synaptic protein and APP mRNA levels. (**C & D**) DN-CREB suppresses baseline levels of synaptic protein and APP protein, and abolishes ApoE3-induced increases of synaptic protein but not APP protein; DP-CREB, in contrast, constitutively increases synaptic protein and APP protein levels (B, representative immunoblots; C, summary graphs of protein levels normalized to Tuj 1 and Ctrl. (EGFP)-ApoE3). NCAD (N-cadherin) and Hsc70 (heat-shock cognate 70) proteins were used as negative controls, and the levels of overexpressed EGFP (used as a molecular manipulation control) or DN-and DP-CREB were also examined as a function of ApoE3 to exclude a possible regulation of the molecular manipulations by ApoE3 itself. Data are means ± SEM (n≥3 independent experiments); statistical significance (*, p<0.05; **, p<0.01; ***, p<0.001) was evaluated with two-way orone-way ANOVA (for APP in all panels, NCAD and Hsc70 in (C)) with Tukey’s multiple comparisons.

To independently confirm these results, we compared in a separate set of experiments the effects of dominant-negative and dominant-positive CREB on synaptic gene expression and on APP both at the RNA (Figure 8B) and protein levels (Figure 8, C and D). Consistent with a central role for CREB in the ApoE-induced stimulation of synaptic gene expression, dominant-negative CREB uniformly decreased synaptic gene expression and blocked the effect of ApoE, whereas dominant-positive CREB constitutively enhanced synaptic gene expression. Again, dominant-negative CREB did not block the effect of ApoE on APP transcription, but decreased overall expression levels. In contrast, dominant-positive CREB (45) constitutively increased APP expression levels (Figure 8, B to D). The levels of two control proteins, N-cadherin and Hsc70, were unaffected. Thus, although cFos is essential only for the ApoE-stimulation of APP expression but not of synapse formation, CREB is likely involved in synapse formation and, to a lesser degree, in *APP* transcription.

How is CREB connected to the MAP kinase pathway that is essential for ApoE-induced synaptic gene activation? To address this question, we monitored CREB phosphorylation as a function of ApoE treatment (Figure 9). CREB phosphorylation in human neurons was potently activated by ApoE with a potency rank order of ApoE4>ApoE3>ApoE2; this activation was ablated by the ApoE receptor blocker RAP (Figure 9, A and B). ApoE-induced CREB phosphorylation was also abolished by the MAP-kinase inhibitor U0126 but not by three other kinase inhibitors: Wortmannin, PKI, or KN93 (Figure 9C). Thus, ApoE binding to its neuronal receptors stimulates MAP kinases, which may phosphorylate CREB to trigger CREB-dependent synaptic gene expression.

**Figure 9.**
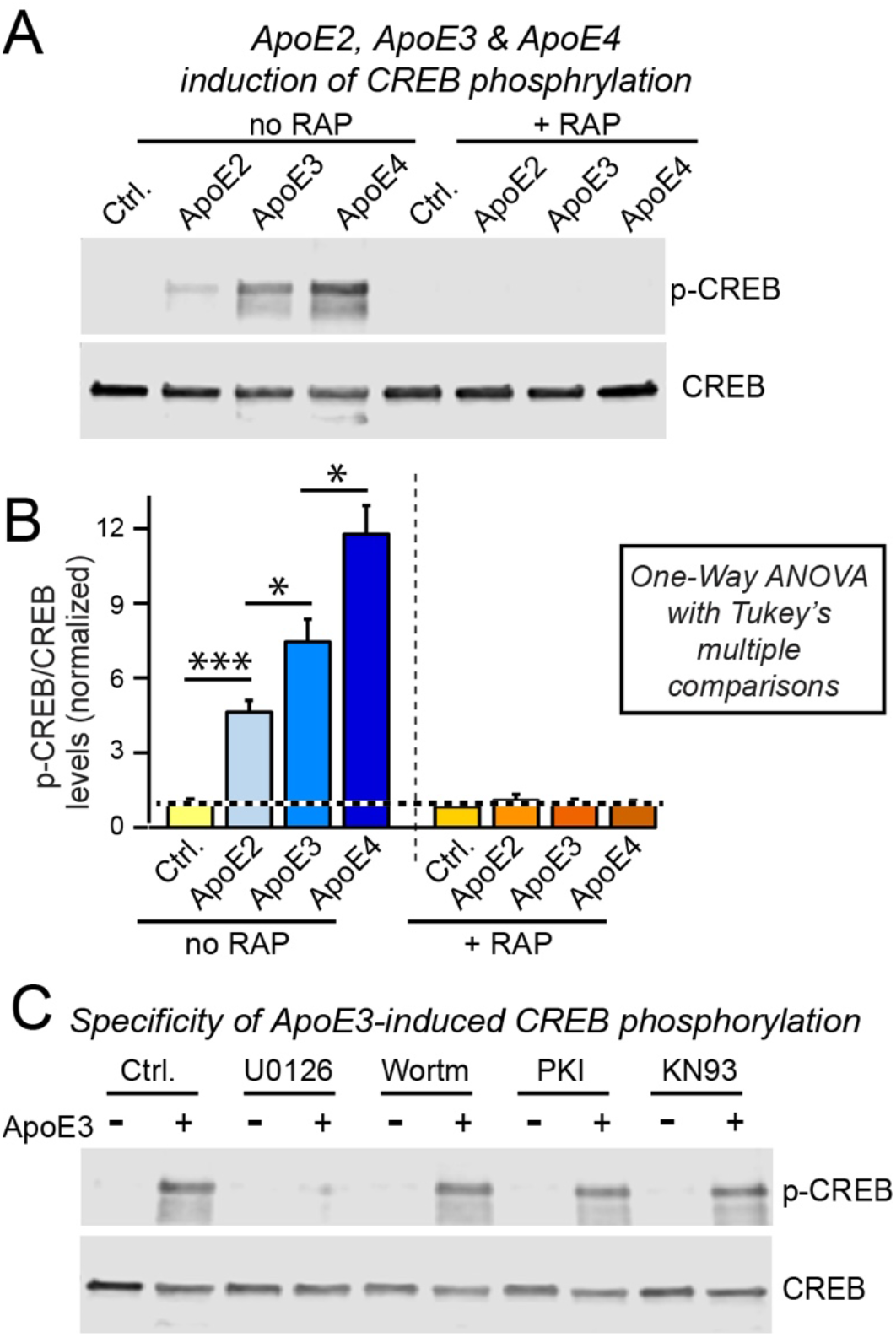
ApoE robustly activates CREB phosphorylation in human neurons grown on MEFs in a MAPK-dependent manner, with ApoE4 being more and ApoE2 being less efficacious than ApoE3. Data are from human neurons cultured on MEFs in the absence of serum; RAP (50 μg/ml), U0126 (50 μM), Wortmannin (0.1 μM), PKI (2 μM) or KN93 (1.0 μM) were added at D10 30 min before recombinant ApoEs (10 μg/ml) or control solutions as indicated, and neurons were analyzed 1 h after ApoE additions. (**A & B**) ApoE activates CREB phosphorylation in human neurons cultured on MEFs with an ApoE4 > ApoE3 > ApoE2 rank potency order, and ApoE-induced CREB phosphorylation is abolished by the ApoE-receptor blocking protein RAP (**A**, representative immunoblots; **B**, summary graphs of phosphorylation levels normalized for controls). Data are means ± SEM (n≥3 independent experiments); statistical significance (*, p<0.05; ***, p<0.001) was evaluated with one-way ANOVA with Tukey’s multiple comparisons. (**C**) The MAP-kinase inhibitor U0126 but not the PI3-kinase inhibitor Wortmannin, the protein kinase A inhibitor PKI, or the CaM kinase inhibitor KN93 blocks the ApoE3-induced phosphorylation of CREB.

### Discussion

In the present paper, we took advantage of a reduced experimental system to investigate the potential signaling role of ApoE specifically in human neurons. We cultured human neurons in isolation from all glial contributions to examine the differential efficacy of ApoE2, ApoE3, and ApoE4 in inducing neuronal signaling. With this approach, we made five major observations (Figure 10). First, we found that ApoE broadly stimulated multiple signal transduction pathways in human neurons in a manner that was inhibited by the ApoE receptor blocking protein RAP; importantly, ApoE2, ApoE3, and ApoE4 exhibited differential signaling efficacies with a potency rank order of ApoE4>ApoE3>ApoE2. Second, we showed that the ApoE-induced stimulation of the MAP-kinase pathway was not neuron-specific, but was also detected in MEFs in which the three ApoE variants also exhibited a potency rank order of ApoE4>ApoE3>ApoE2. Third, we found that a major downstream effect of ApoE signaling in neurons was stimulation of synapse formation that increased the number of synapses without changing the size of the neurons or their dendrites. This ApoE-induced stimulation of synapse formation involved increased expression of synaptic genes, suggesting that it included gene transcription as a basic mechanism, similar to the ApoE-induced increase in APP and Aβ synthesis we have previously described (37). Fourth, we showed that the ApoE-induced increase in synapse formation required MAP-kinase activation similar to ApoE-induced APP transcription, suggesting that the ApoE-stimulated MAP-kinase pathway has multiple readouts. Fifth and finally, we documented that, unlike the ApoE-induced increase in APP transcription that required cFos, the ApoE-induced increase in synapse formation was independent of cFos but required another transcription factor, CREB.

**Figure 10.**
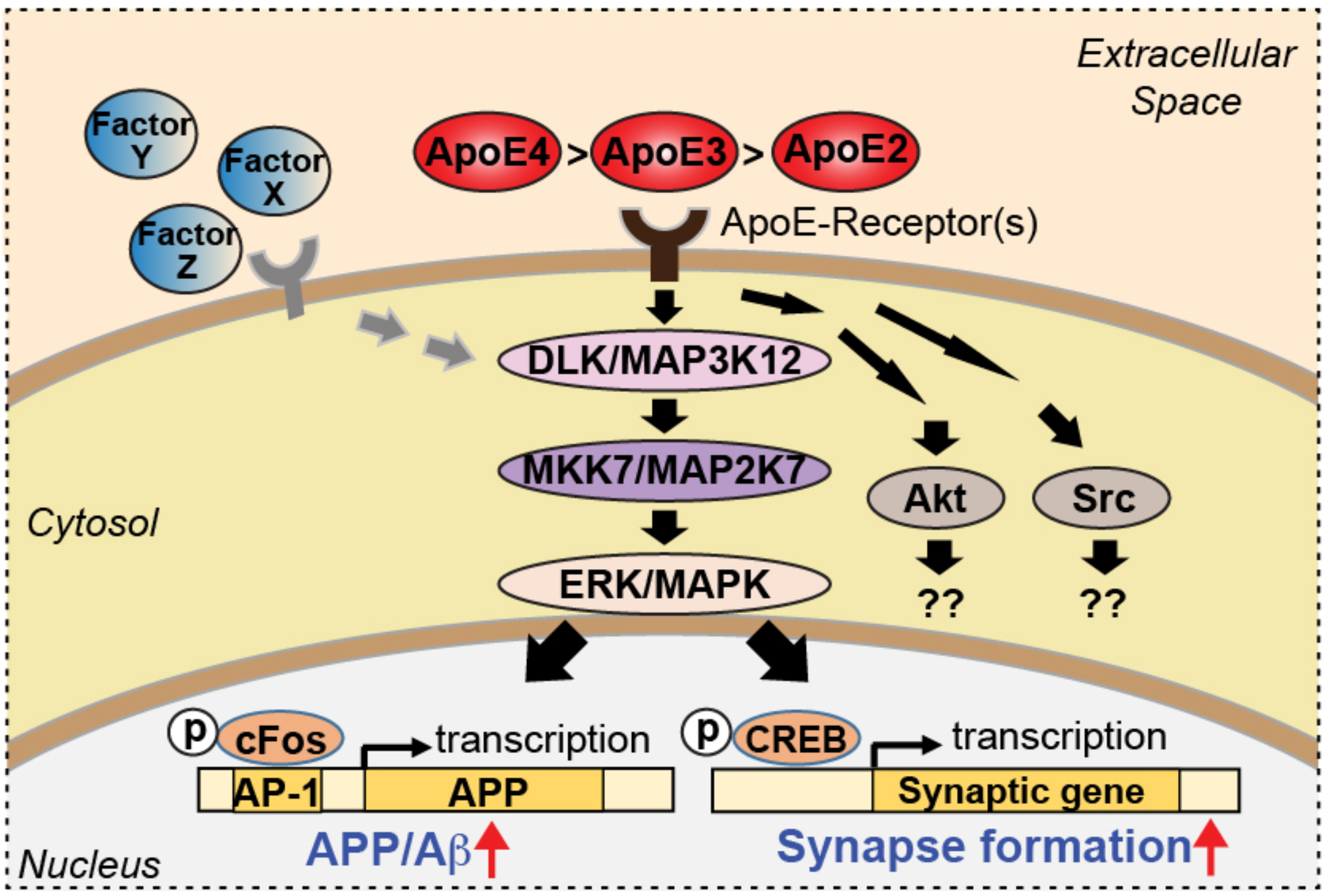
Summary cartoon of differential neuronal signaling mediated by ApoE2, ApoE3, and ApoE4. ApoE-binding to neuronal receptors activates multiple parallel signaling pathways, including a MAP-kinase cascade that induces both cFos-and CREB-phosphorylation and stimulates APP transcription and synaptogenesis.

Viewed together with earlier results (37), these findings suggest that in brain, one of the biological functions of ApoE is as a signaling molecule that activates diverse pathways in neurons (Figure 10). ApoE broadly yet selectively induced multiple pathways. For example, we showed that ApoE stimulates synapse formation but not dendritic arborization, and we found earlier that ApoE stimulates transcription of *APP* but not of the related genes *APLP1* and *APLP2* (37). These results are compatible with other data suggesting a major signaling role of ApoE in microglia (reviewed in (46)), and support with the notion that activation of astrocytes and/or microglia enhances ApoE secretion, which then in turn acts as a signaling molecule to produce diverse effects on both microglia and on neurons. We would like to argue that the proposed signaling role of ApoE could serve as a repair signal and injury response, with increased synaptogenesis serving to compensate for potential losses of synaptic function during pathological processes.

As always, our conclusions are contingent on the specific conditions of our experiments, which involved human neurons cultured on MEFs or a non-cellular substrate (matrigel) in the absence of glia or serum. When human neurons are cultured in the presence of glia that are activated by the culture conditions and secrete an array of abundant factors, the application of additional ApoE has no specific effect, most likely because the glial factors occlude the effect of ApoE (Figure 3 and Supplemental Figure 6C)(37). The signaling events defined in our experiments thus represent processes that are normally obscured unless under a glia-free condition, which can be by definition regarded as non-physiological. It is important to note, however, that all studies in which cultured neurons and glia are removed from their normal three-dimensional organization are likewise non-physiological. After all, in any experimental condition utilizing cultured neurons, the normal cellular interactions are disrupted, both glia and neurons exist in an abnormal state, and many properties – such as LTP in neurons – are lost. Nevertheless, such reduced systems are valuable and informative. Our specific approach provides high clarity by excluding glial signals, and, as a result, allows defining neuronal responses that would otherwise be blunted or concealed under conventional settings but are likely to be physiologically operating and important in a normal cellular context. This highly defined approach is akin to a biochemical reconstitution experiment of a protein complex, in which the normal cellular complexity is purposely eliminated to allow reconstruction of a specific molecular function.

With these considerations in mind, we posit that the signaling pathways we define here provide insight into fundamental properties of ApoE and its receptors in neurons, regardless of whether this is a major or minor contributor to human brain function *in vivo.* Our most important finding here is that ApoE binding to its receptors on neurons induces a broad activation of multiple signaling pathways that operate via multiple mechanisms and impact selective downstream processes. Although both synapse formation and APP transcription were induced by ApoE in a manner requiring activation of MAP kinases, the downstream effectors were different in that the former required CREB whereas the latter involved cFos. In all of the recorded ApoE activities, ApoE variants exhibited differential efficacies with a potency rank order of ApoE4>ApoE3>ApoE2. In these activities, ApoE acted by binding to classical ApoE receptors because its actions were inhibited by RAP specifically. Owing to our reductionist design, these findings were clearly not confounded by signaling of TREM2, a reported alternative ApoE receptor implicated in AD, because TREM2 is not detectably expressed in MEFs or human neurons and, as a non-canonical putative ApoE receptor, TREM2 is not expected to bind to RAP. Neurons express multiple classical ApoE receptors, most prominently LRP1, and the precise ApoE receptor(s) involved in the processes we study remains unknown.

How do the present observations relate to the contribution of allelic ApoE variants to AD pathogenesis? It seems counterintuitive that ApoE4, which predisposes to AD and thus presumably to neurodegenerative synapse loss, should be more effective in promoting synapse formation than ApoE2, which protects against AD and synapse loss. However, ApoE is not required for overall normal brain function as evidenced by the generally normal cognitive performance of ApoE-deficient human patients (12–14). This fact suggests that the signaling function of ApoE does not contribute an essential component to brain performance under non-pathological conditions and is not required for normal synapse formation. If ApoE signaling kicks in under pathophysiological conditions, the increased efficacy of ApoE4 may not be functionally relevant for individual pathophysiological events, but cause differential effects on neuronal survival over prolonged time periods. The fact that the genetic ApoE variants show such robust and consistent differences in their respective potency of acting these neuronal signaling pathways seems to us to be more important than the precise nature of this signaling, because this fact provides a potential explanation for why ApoE4 is deleterious and ApoE2 is protective in AD pathogenesis, even if the specific nature of AD pathogenesis and the role of ApoE-signaling in it remain undefined.

The finding that ApoE2, ApoE3, and ApoE4 exhibit differential efficacies in neuronal signaling that mirrors their ApoE-receptor binding properties has potential therapeutic implications. If ApoE-induced neuronal signaling is involved in AD pathogenesis, decreasing such signaling that is enhanced in ApoE4 carriers would be a viable pathway for therapy. Achieving such a therapy, however, requires not only validating the proposed mechanism of the role of ApoE4 in AD pathogenesis – something that has arguably not yet been achieved for any hypothesis – but also identifying the specific receptor involved in neuronal ApoE signaling. Multiple ApoE receptors are expressed in neurons. Which of these receptors mediates the neuronal ApoE signaling characterized here, however, is unknown. Identification of this receptor is thus the most important next goal towards a development of pharmacological agents that could specifically dampen ApoE4 signaling and thereby ameliorating the increased risk for AD in ApoE4-expressing population. Accomplishing this goal will be a primary focus in forthcoming studies.

## Materials and Methods

### Study Design

No statistical methods were used to predetermine sample size. No samples were excluded from analysis. Two major sets of experiments (Figure 1 Supplemental Figure 3; Figure. 5 and Supplemental Figure 4) were randomized and investigators were blinded to sample identities during experimentation and outcome assessment. All experiments involving mice and human embryonic stem (ES) cells were performed in accordance with Stanford University and federal guidelines with approval of appropriate protocols by the various regulatory committees.

### Culture of principal cell types

The cell culture procedures were performed as reported previously (37), with details described in Supplementary Materials. H1 human ESCs were obtained from WiCell Research Resources (Wicell, WI) and maintained in feeder-free condition. Mouse glia were cultured from the cortex of newborn CD1 mice. Murine embryonic fibroblasts (MEFs) were isolated from mouse embryos of CF-1 strain (Harlan Laboratories, Inc.) harvested at 12.5-13.5 postcoitum (p.c.).

### Generation of human neurons from H1 human ES cells

Human neurons were generated from H1 cells essentially as described (37). In short, ES cells were detached with Accutase and plated onto matrigel-coated 6-well plates (4×104 cells/well) on day −2. Lentiviruses expressing Ngn2 and rtTA were prepared as described below, diluted in fresh mTeSR1 medium, and added to the ESCs on day −1. Doxycycline (2 mg/l, to activate Ngn2 expression) was added on day 0 (D0) in DMEM/F-12 medium with N2 supplement without morphogens. Puromycin (1 mg/l) was added on D1 in fresh DMEM-F12/N2 + doxycycline medium for selection up to 48 hours. On D3, differentiating neurons were detached with Accutase and re-plated on cultured mouse glia, MEFs, or just matrigel-coated 24-well plates (2×10^5^ cells/well), and maintained in NBA/B-27 medium with no doxycycline. Lentiviral infection of iN cells were performed on D4 as described below; ApoE incubations were initiated on D10 and lasted until neurons were analyzed for various parameters. For mRNA and synaptic protein measurements, the assays were performed on D12 after ApoE treatments of 2 days, unless otherwise specified in time course studies. For immunoblotting analysis of protein phosphorylation, the ApoE was administered for 2 hours, and neuronal cultures were harvested immediately. For synaptic density and morphology analysis, neurons were fixed for immunofluorescence on D16 or D23-35. The electrophysiological recordings were done during D23-25.

### Production of recombinant proteins

Recombinant ApoE2, ApoE3, and ApoE4 were produced in HEK293 (FreeStyle 293-F) cells (ThermoFisher) or bacteria (*E. coli* BL21 strain) essentially as described previously (37). The detailed procedures are provide in Supplementary Materials. Recombinant proteins of glia-secreted factors proteins for screening experiments (Supplemental Figure 6C) were produced from HEK293T cells transfected with plasmids encoding human proteins by a standard calcium phosphate protocol. For detailed procedure and cDNA information please see our previous report (37). For production of recombinant RAP, pGEX-KG-RAP (provided by Dr. Joachim Herz, UT Southwestern Medical Center) was expressed in BL21 bacteria, and GST-RAP was purified as described above, except that thrombin cleavage (10 U per mg protein, overnight at 4°C) was used instead of rhinovirus 3C protease as a final step.

### Lentivirus-mediated gene expression

Lentiviruses were produced in HEK293T cells as described (47), from the following plasmids: i. lentiviruses to trans-differentiate ES cells into human neurons: TetO-Ngn2-P2A-puromycin and rtTA (47); ii. lentiviruses for overexpression of DLK, MKK7 and MBIP: pLX304-DLK, clone ID: HsCD00413295; pLX304-MBIP, HsCD00420627; pCW45-MKK7, HsCD00298961 (Harvard Medical School); iii. DN-cFos (48), DN-CERB (43), DN-MEF2A (44) and DP-CREB (45): with synthesized human cDNA fragments modified from validated rodent sequences cloned into AgeI and EcoRI sites on lentiviral vector FUGW (Addgene plasmid # 14883)

### Suppression of gene expression using RNAi or CRISPR/Cas9

i. RNAi of DLK. shRNA to DLK (sequence: ACTCGTATTCCTTGTACATAG, TRC number: TRCN0000231658) and control shRNA (sequence: TAAGGCTATGAAGAGATAC; SHC016) were purchased from Sigma-Aldrich in lentiviral vector pLKO.1-puro. The DLK shRNA targets the 3’UTR of DLK mRNA and does not affect expression of rescue DLK, while the control shRNA contains at least four mismatches to any human or mouse gene and was demonstrated by the manufacturer to target zero gene using microarray analyses; ii. MKK7 CRISPR. Lentiviral CRISPR/Cas9-mediated inhibition of human MKK7 expression was performed using a plasmid (lentiCRISPR v2; Addgene plasmid # 52961) that co-expresses Cas9 nuclease with a single guide RNA (sgRNA). The MKK7 sgRNA (sequence: GCTTCAGCTTTGCTTCCAGG) targets exon 1 with a cleavage site at amino acid 13 and was designed using web-based tools. The control sgRNA targets EGFP (EGFP sgRNA4; sequence: GGAGCGCACCATCTTCTTCA; Addgene plasmid # 51763) and was cloned into the same Cas9-expressing vector. The efficacy of the inhibition of gene expression by RNAi, or CRISPR/Cas9 was assessed by qRT-PCR as reported previously (37).

### Immunofluorescence labeling experiments

Immunofluorescence staining experiments and image acquisition and analyses were performed essentially as described (37). Briefly, cultured neurons were fixed in 4% paraformaldehyde, 4% sucrose in PBS, permeabilized with 0.2% Triton X-100 in PBS, and blocked with 5% goat serum in PBS. Cells were incubated with primary antibodies diluted in blocking buffer overnight at 4°C, washed 3 times, and incubated with secondary antibodies in blocking buffer for 1 h at room temperature. Samples were then mounted on glass slides for confocal imaging. The following antibodies were used: mouse anti-MAP2 (Sigma M1406, 1:1,000), rabbit anti-synapsin (E028, 1:1,000), rabbit anti-HOMER1 (Synaptic Systems 160003; 1:1000), mouse anti-Tuj1 (Covance MMS-435P-250, 1:2,000); Alexa-488-, Alexa-546-, and Alexa-633-conjugated secondary antibodies (Invitrogen). Immunofluorescence signals were visualized using a Nikon A1 confocal microscope with constant image settings. Neurons were randomly chosen in confocal images. Synapsin-positive or Homer1-positive synaptic puncta were quantified for puncta density per dendritic length, size, and intensity. Total dendritic length and cell body size were quantified based on tracing of MAP2 signals.

### Immunoblotting and protein quantifications

Neurons and cells were lysed in RIPA buffer (50 mM Tris-HCl pH 8.0, 150 mM NaCl, 0.1% SDS, 0.5% sodium deoxycholate, 1% Triton-X 100, plus a cocktail of protease inhibitors (Roche)), and lysates were analyzed by SDS-PAGE in the presence of DTT (0.1 M). Immunoblotting and quantitative analysis were performed with fluorescent-labeled secondary antibodies and an Odyssey Infrared Imager CLX and software Image Studio 5.2.5 (LI-COR Biosciences). Signals were normalized for neuronal TUBB3 probed on the same blots as loading controls. Antibodies used were as follows: GluA1 (Abcam ab1504, 1:500), PSD-95 (L667, 1:1,000), synapsin (E028, 1:2,000), synaptophysin (K831, 1:1,000), synaptotagmin-1 (V216, 1:1,000), synaptobrevin-1(T2797, 1:1,000), Tuj1 (Covance MMS-435P-250, 1:2,000), APP (Millipore MABN380, 1:2,000), DLK (Sigma SAB2700169, 1:1,000), phospho-MKK7 Ser271/Thr275 (Cell Signaling 4171, 1:500), MKK7 (Santa Cruz sc-25288, 1:500), phosphor-ERK1/2 Thr202/Tyr204 (Cell Signaling 9106, 1:1,000), ERK1/2, (Cell Signaling 4695, 1:1,000), phospho-Akt Ser473 (Cell Signaling 9271, 1:1,000), Akt (Cell Signaling 2966, 1:1,000), phosphor-Src Tyr 418 (ThermoFisher 44-660G, 1:500), Src (ThermoFisher AHO1152, 1:1,000) phospho-JNK Thr183/Tyr185 (Cell Signaling 9255, 1:250), JNK (Cell Signaling 9252, 1:500), phospho-c-Fos Ser374 (Santa Cruz sc-81485, 1:500), phosphor-CREB (Cell Signaling 9198, 1:1,000), CREB (Cell Signaling 9104, 1:1,000), β-actin (Sigma a1978, 1:1,000); ApoE (ThermoFisher 701241, 1:1,000)

### Gene Expression Analyses

To determine the mRNA levels of genes of interest in cultured cells, quantitative RT-PCR (qRT-PCR) measurements were performed on total RNA (isolated with PrepEase RNA Spin Kit, Affymetrix) using TaqMan probes with VeriQuest Probe One-Step qRT-PCR Master Mix (Affymetrix) and an Applied Biosystems 7900HT apparatus. The pre-designed TaqMan primer/probe sets were purchased from Integrated DNA Technologies and tested to show no or minimal cross-species reactivity in pure human neuronal and mouse glial cultures (37). MAP2 and GAPDH were used as endogenous reference genes. The assay IDs of all TaqMan primer/probe sets used are: human MAP2, Hs.PT.58.20680759; human GAPDH, Hs.PT.58.40035104; human APP, Hs.PT.56a.38768352; human SYN1, Hs.PT.58.4027324; human SYP, Hs.PT.58.27207712; human SYT1, Hs.PT.58.19615550; human SYB1/VAMP1, Hs.PT.58.23319147; human PSD95/DLG4, Hs.PT.58.20575145; human GluA1/GRIA1, Hs.PT.58.40318075; mouse Gapdh, 4352932-0809025; mouse Apoe, Mm.PT.58.33516165.

### Electrophysiology

Whole-cell voltage clamp recordings were performed at room temperature on human neurons at D23-25, with 3-3.5 MΩ borosilicate patch pipettes filled with an internal solution containing (in mM) the following: 135 CsMeSO3, 8 NaCl, 10 HEPES, 0.25 EGTA, 4 MgATP, 0.3, Na3GTP, 5 Na-phosphocreatine, 2 MgCl2, and 2 QX314 (pH adjusted to 7.30 with CsOH). Cells were held at −70mV in the bath solution containing (in mM): 140 NaCl, 10 HEPES, 10 glucose, 5 KCl, 3 CaCl2 and1 MgCl2 (pH adjusted to 7.40 with NaOH). For mEPSC recordings, TTX (1 μM) and picrotoxin (50 μM) were added to the bath solution. All electrophysiological recordings were performed with Multiclamp 700B amplifiers (Molecular Devices) and analyzed using Clampfit 10.4 (Molecular Devices).

### Statistics

Quantitative data are presented as means ± SEM. All experiments were independently repeated at least 3 times. Statistical analyses were conducted using Prism 6 software (GraphPad Software, Inc.). Statistical comparisons between groups were analyzed for significance by one-way or twoway analysis of variance (ANOVA) with Tukey’s post-hoc test. Our data meet the normal distribution assumption of the tests. There is an estimate of variation within each group of data, and the variance is similar between the groups that are being statistically compared.

## AUTHOR CONTRIBUTIONS

Y.A.H. and T.C.S. planned all experiments, analyzed the data, and wrote the paper; Y.A.H., B.Z. and A.M.N. performed all experiments; and M.W. provided advice on data analyses and the manuscript.

## ACKNOWLEDGEMENTS

We thank Dr. J. Herz for the RAP expression vector. This work was supported by NIH grants RF1 AG04813101 (to T.C.S. and M.W.) and K99 AG054616 (to Y.A.H.). M.W. is a Tashia and John Morgridge Faculty Scholar from the Child Health Research Institute at Stanford and a Howard Hughes Medical Institute Faculty Scholar.

## Supplemental Data

### Materials and Methods

#### Culture of principal cell types

a. H1 human ESCs were obtained from WiCell Research Resources (Wicell, WI), maintained in mTeSR1 medium (Stem Cell Technologies) without feeder cells, and used at intermediate passage numbers (~50-60) to generate human neurons, as described previously (47). Mycoplasma testing and karyotyping were performed regularly (approximately every 10 passages). The H1 ES cell line is not listed in the database of commonly-misidentified cell lines maintained by ICLAC.
b. Mouse glia were cultured from the cortex of newborn CD1 mice. The cortex was dissected, digested with papain (10 U/ml) for 20 minutes at 37^°^C, and harshly triturated. Cells were plated onto T75 flasks in Dulbecco’s Modified Eagle Medium (DMEM) supplemented with 10% fetal bovine serum (FBS). At 80-90% confluence, glial cultures were trypsinized and re-plated at lower density (20-30% confluence) in DMEM + 10% FBS. This re-plating procedure was repeated 2-3 times (typically 3–4 days apart) to remove mouse neurons before cultured glia were used for analysis or co-culture experiments with human neurons. No antibiotics or other drugs were used in glial cultures.
c. Murine embryonic fibroblasts (MEFs) were isolated from mouse embryos of CF-1 strain (Harlan Laboratories, Inc.) harvested at 12.5-13.5 postcoitum (p.c.). Briefly, embryos were dissected out of terminally-anesthetized mice. The head and internal organs were removed, and the remaining carcasses were finely minced, trypsinized into single-cell suspensions, and plated onto T75 flasks. The cultured MEFs (P0) were frozen or briefly expanded (a maximum of 3 times, P3) before they were used for experiments.

#### Production of recombinant ApoE proteins

Recombinant ApoE2, ApoE3, and ApoE4 were produced in HEK293 (FreeStyle 293-F) cells (ThermoFisher) or bacteria (*E. coli* BL21 strain). Expression vectors for human ApoE2, ApoE3, and ApoE4 were constructed using ApoE3 cDNA in a pDONR221 vector (Gateway recombination system) obtained from Harvard Medical School (Harvard PlasmID Database, clone ID: HsCD00044600). Site-directed mutagenesis was used to generate ApoE2 (C526T) and ApoE4 (T388C) cDNAs. The ApoE2, ApoE3, and ApoE4 cDNAs were cloned into the lentiviral vector pLX304 (Addgene plasmid # 25890) using Gateway LR Clonase II (Invitrogen); the control plasmid was FUGW (Addgene plasmid # 14883), used to monitor transfection efficiency. 293-F cells were cultured in suspension in serum-free FreeStyle 293 Expression Medium (ThermoFisher, protein-free and chemically defined formulation) and transfected with control or ApoE expression plasmids using lipid-based FreeStyle MAX Reagent (ThermoFisher) following the manufacturer’s instructions. Supernatants from transfected 293-F cells were harvested 6 days after transfection and assessed by SDS-PAGE to determine the purity and yields of recombinant ApoE proteins (Supplemental Figure 5A). The ApoE supernatants were diluted to proper concentration by fresh NBA/B27 (typically 50-80X) and added into cultured neurons for ApoE stimulation. The supernatant from control- (FUGW-) transfected 293-F cells was diluted the same way and added onto neurons as the control condition, in order to control for effect of other secreted proteins. HEK293 ApoE proteins were used in all ApoE stimulation experiments unless otherwise indicated. In the E. coli BL21 strain, ApoE proteins were expressed as GST fusion proteins in modified pGEX-KG vectors harboring in multiple cloning site (*BamHI-EcoRI*) a cleavage site recognized by human rhinovirus 3C protease (PreScission Protease): LEVLFQ/GP (DNA sequence: ctggaagttctgttccaggggccc). The cDNAs encoding ApoE2, ApoE3, and ApoE4 (mature, 299 amino acids) for the HEK293 cell expression experiments were cloned into BamHI and HindIII sites. GST-ApoE proteins were purified essentially as described (49). Bacteria were grown to OD600 of 0.5 and protein expression was induced with Isopropyl β-D-1-thiogalactopyranoside (IPTG, 0.05 mM) for 6 h at room temperature. Bacteria were then pelleted and resuspended in solubilization buffer (0.5 mg/ml lysozyme in PBS, 1 mM PMSF, 1 mM EDTA, and an EDTA-free protease inhibitor cocktail). Cell were lysed by liquid nitrogen freeze-thaw cycling and sonication. After centrifugation to remove insoluble components of cell lysates, proteins were affinity-purified with glutathione sepharose beads and collected by cleavage with rhinovirus 3C protease (PBS, 16 hours at 4 °C).

---

**Supplemental Figure 1.**
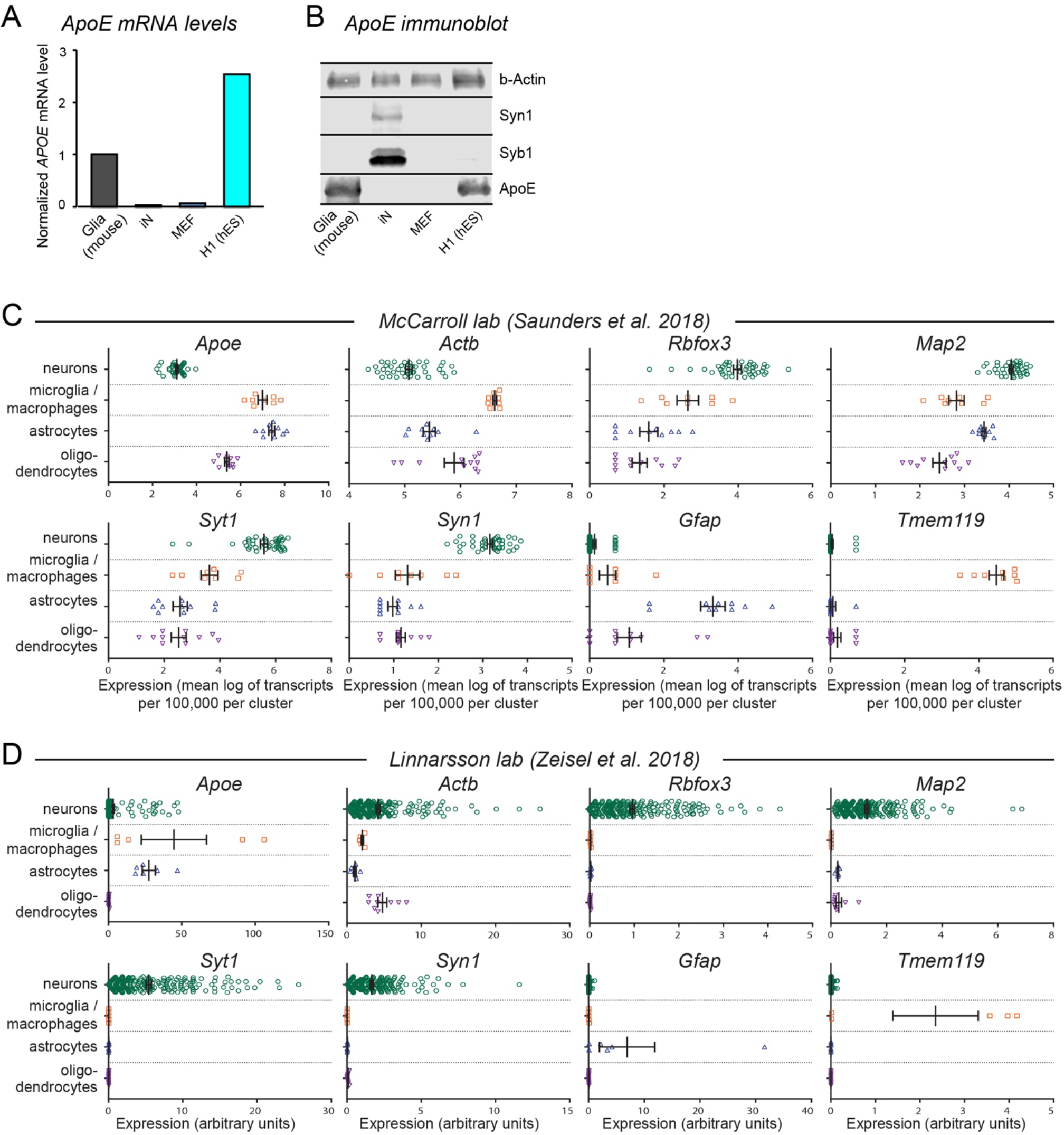
ApoE mRNA and protein is highly expressed in human ES cells but not in human neurons derived from the ES cells or in MEFs (A & B), while analyses of published single-cell RNAseq datasets reveal that ApoE is expressed at only low levels in neurons, but high levels in astrocytes and microglia (C & D) (**A**) ApoE mRNA levels are high in mouse glia and human H1 ES cells, but barely detectable in human neurons (iN) and mouse embryonic fibroblasts (MEFs) as determined by quantitative RT-PCR using species-specific primers. Levels were normalized to GAPDH as an internal standard. Human neurons were examined at day 10 after induction (D10) with neurons cultured on matrigel in the absence of glia, MEFs, or serum. (**B**) Immunoblotting detects robust levels of ApoE protein in cultured mouse glia and human H1 ES cells, but not in human neurons (iN) or MEFs. β-actin was analyzed as an internal standard, and synapsin-1 (Syn1) and synaptobrevin-1 (Syb1) as neuron-specific synaptic proteins. (**C & D**) Single-cell RNAseq data obtained by Saunders et al. (**C**)(39) and Zeisel et al. (**D**)(40) were analyzed for relative expression levels of the indicated genes. Using the clusters as defined by each of these studies, cell types were grouped into the following populations: neurons, microglia/macrophages, astrocytes, and oligodendrocytes. Gene expression levels were compiled from each dataset with no further processing, using the units as displayed from each database.

**Supplemental Figure 2.**
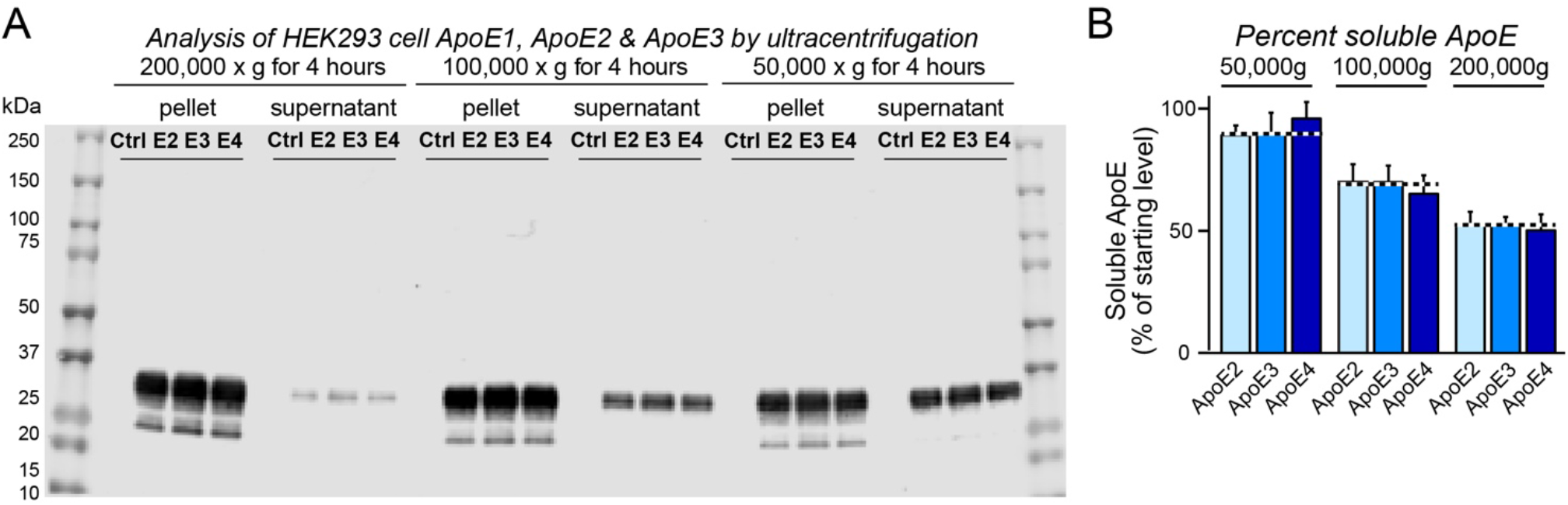
Ultracentrifugation analysis of ApoE2, ApoE3, and ApoE4 produced in transfected HEK293 cells fails to detect differences in physical states. (**A & B**) ApoE2, ApoE3, and ApoE4 harvested from the medium of transfected HEK293 cells are pelleted by similarly high ultracentrifugation forces, suggesting that they are components of similarly sized particles and not large aggregates. ApoE2, ApoE3, and ApoE4 as well as control solutions were subjected to ultracentrifugation at the indicated g-forces for 4 hours, and the resulting supernatants and pellets were analyzed by quantitative immunoblotting using fluorescent secondary antibodies. (**A**),representative immunoblot; (**B**), summary graph of soluble ApoE protein shown as means ± SEM (n≥3 independent experiments), with no statistical significance by one-way ANOVA with Tukey’s multiple comparisons.

**Supplemental Figure 3.**
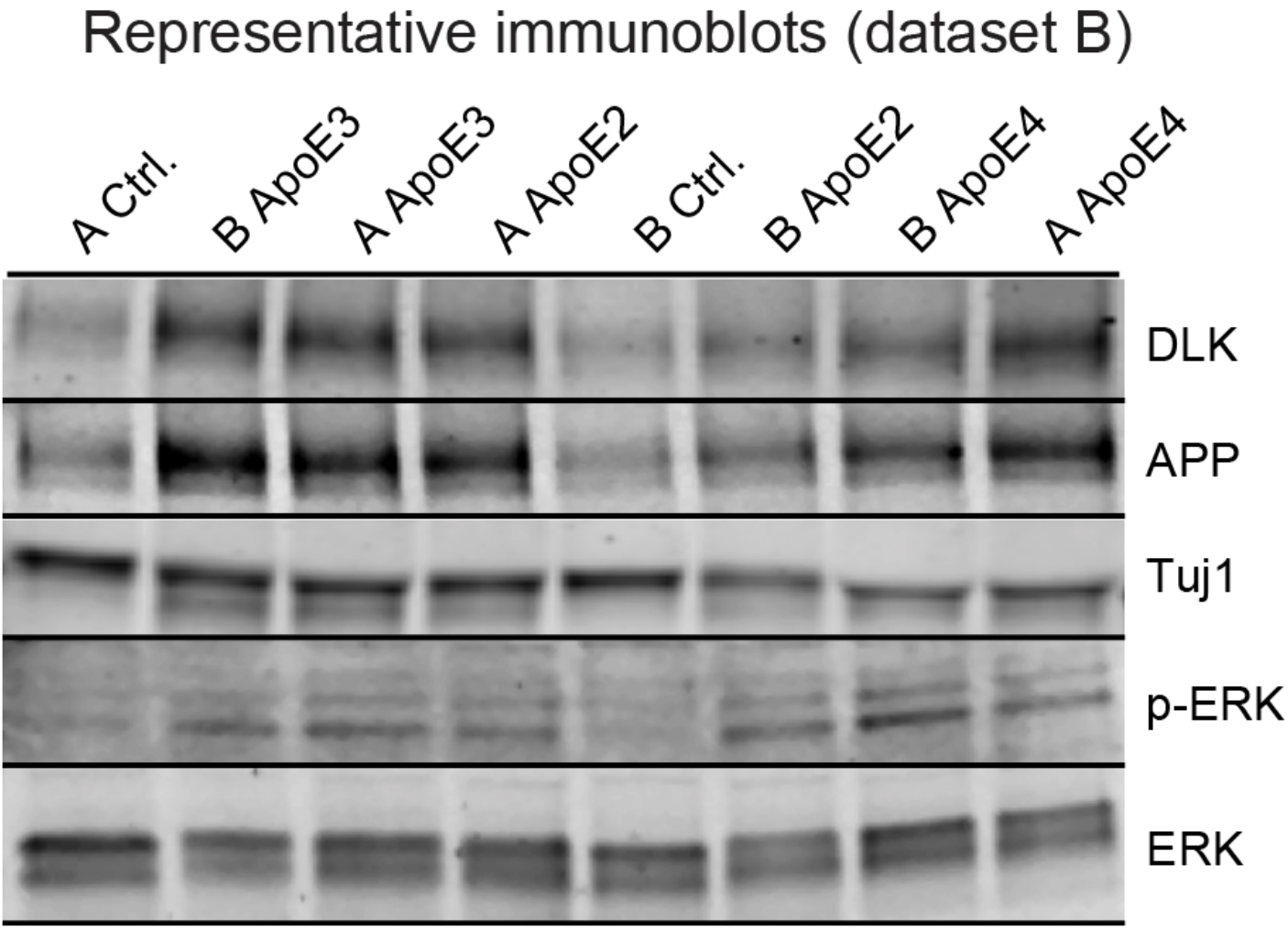
Representative immunoblot for dataset B in the independent analyses of the reproducibility of ApoE-induced neuronal signaling shown in Figure 1. Representative immunoblots from dataset B, performed by one of two independent experimenter notated as A and B. For summary graphs and experimental conditions, see Figure 1. In blinding experimenters to the nature of the reagents as much as possible, experiments were performed such that recombinant ApoE proteins, as well as a control preparation of supernatant from GFP-transfected cells, were harvested from independently transfected FreeStyle HEK 293 cells and diluted to the final concentration in separate preparations by experimenters A and B. These two sets of solutions were then passed to a third researcher, who randomly coded the samples and distributed them back to experimenter A and B. Each experimenter grew separate cultures and began treatment with the randomly coded ApoE solutions at D10 for 48 hours. After ApoE treatment, cells were harvested and were then subjected to immunoblot (Figure 1, A and B; Supplemental Figure 3) or qRT-PCR analyses (Figure 1C) to probe for expression of the indicated proteins and genes. Following the completion of all data analysis, the sample identities were decoded and compared. This parallel, blinded testing was repeated three times. Labeling of the immunoblot depicted here and in Figure 1 shows sample identity following de-coding of the data.

**Supplemental Figure 4.**
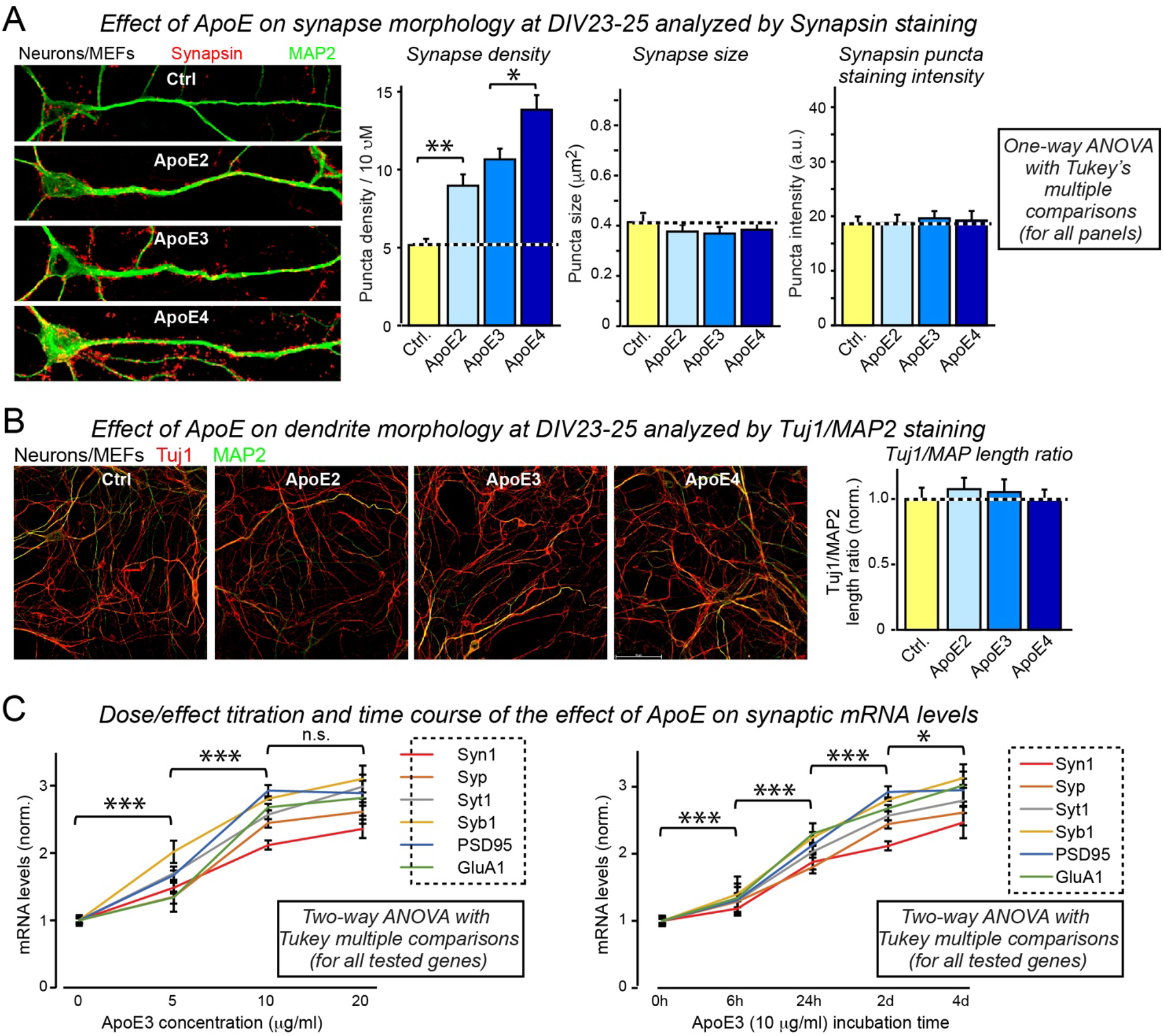
ApoE stimulates synaptogenesis without affecting neuronal dendritic morphology also at later stages of cultured human neurons (A & B), and ApoE3-induced increases in synaptic protein mRNA are dose-and time-dependent (C). (**A**) ApoE’s synaptogenic effect was examined at a later culture time (D23-25) corresponding to the time point of the electrophysiological recordings in Figure 5 (left, representative images of human neurons cultured on MEFs, treated with ApoE from D10 and fixed at D23-25 for immunostaining to label dendrites by MAP2 fluorescence signals and to label synapses by synapsin; right, summary graphs of synapse density, calculated by synapsin puncta number over the length of dendrite, synapse size and synaptic synapsin-staining intensity). (**B**) ApoE does not change neurite morphology even after prolonged culture, as assayed by confocal imaging (left, representative images of human neurons cultured on MEFs, prepared as described in (**A**) but stained for Tuj1 and MAP2; right, summary graph of the length ratio of Tuj1-and MAP2-positive processes). (**C**) Measurements of the ApoE3-concentration-dependence of the induction of synaptic protein mRNA expression (left) and of the time course of ApoE3-induced synaptic protein mRNA expression (right; at 10 μg/ml ApoE3). Human neurons were cultured on MEFs and treated with ApoE3 at days D10 to D12 after neuronal induction for the dose-response curve, or at the indicated times before D12 for the time course. All qRT-PCR assays were performed at D12 and detected only human transcripts; MAP2 was used as an internal control that relates the synaptic protein mRNA levels to those of an internal neuronal marker to correct for the increasing degree of neuronal differentiation during the course of the experiment. mRNA levels were normalized to the untreated control or the 0 h time point. Data are means ± SEM (n≥3 independent experiments); statistical significance (*, p<0.05; **, p<0.01; ***, p<0.001) was evaluated with two-way (**A**) or one-way ANOVA (**B** and **C**) with Tukey’s multiple comparisons.

**Supplemental Figure 5.**
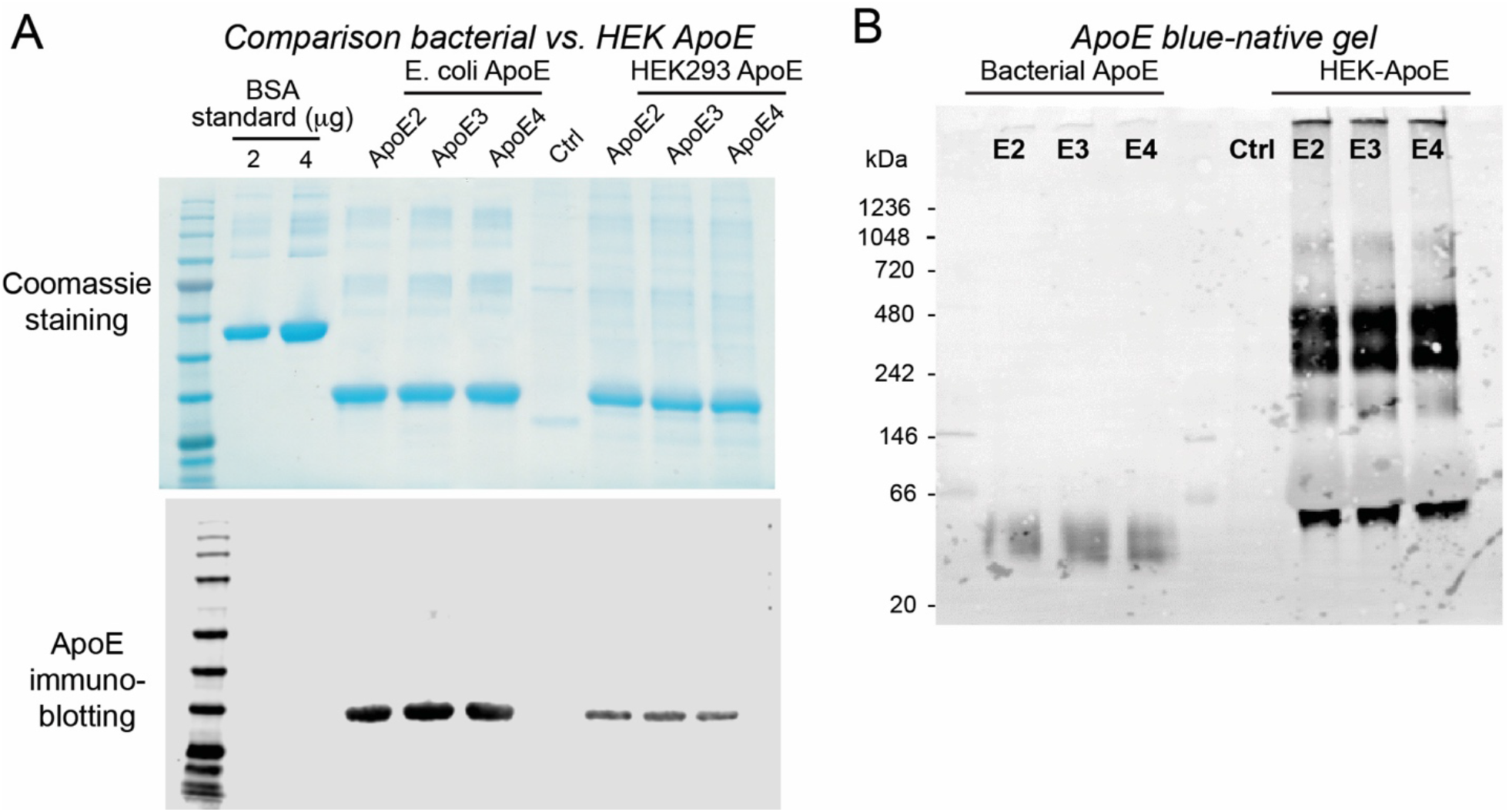
Comparative analysis of ApoE2, ApoE3, and ApoE4 produced in E. coli and in transfected HEK293 cells using denaturing SDS-PAGE (A) and blue-native SDS-PAGE (B) (**A**) SDS-PAGE analysis of recombinant ApoE2, ApoE3, and ApoE4 proteins produced in transfected HEK293 cells and in bacteria (*E. coli).* Left, Coomassie Blue staining; right, immunoblotting for ApoE. Bovine serum albumin (BSA) was analyzed on the same gel at different concentrations as a protein standard to calculate ApoE concentrations. (**B**) Analysis of ApoE2, ApoE3 and ApoE4 proteins produced in bacteria (*E coli)* and HEK293 cells by non-denaturing electrophoresis followed by immunoblotting for ApoE. This blue native polyacrylamide gel electrophoresis (BN PAGE) technique provides a near-neutral operating pH and detergent compatibility for molecular weight estimations. No isoform-specific difference was observed in both bacterial and HEK293 ApoEs, but the apparent size of ApoE particles was dramatically different, suggesting a different physical state.

**Supplemental Figure 6.**
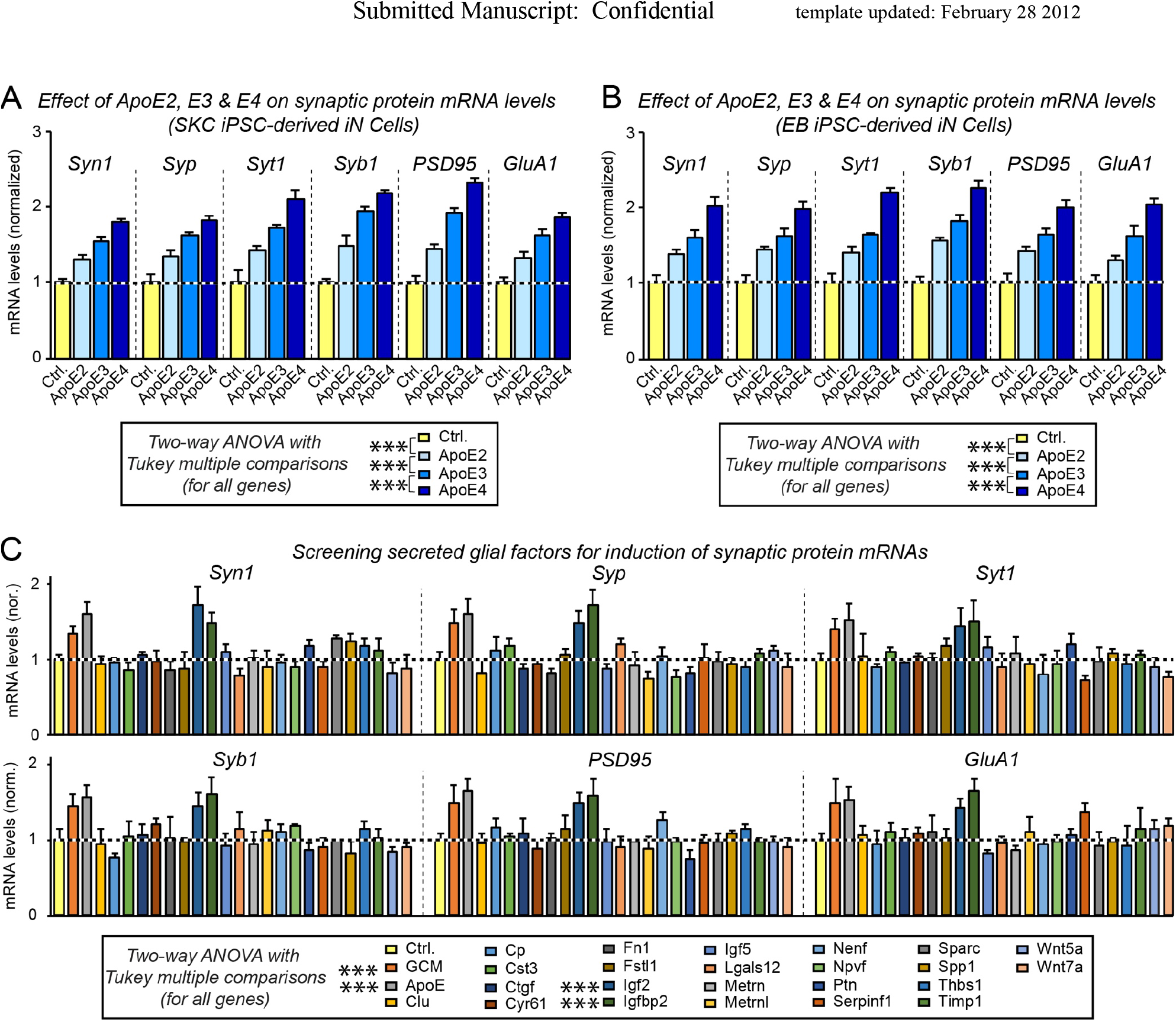
ApoE2, ApoE3, and ApoE4 also stimulate synaptic gene transcription with a rank-potency order of ApoE4>ApoE3>ApoE2 in human neurons derived from two different iPS cell lines (A & B), and other factors secreted by glia in addition to ApoE also stimulate synaptic gene transcription (C) (**A & B**) ApoE2, ApoE3 and ApoE4 differentially increase mRNA levels of synaptic genes in human neurons that were derived from two different iPSC lines (**A**, SKC; **B**, EB) and cultured on MEFs. Experiments were carried out as described in Figure 3. (**C**) Screening 24 secreted proteins that are abundantly produced by cultured mouse glia reveals three factors (ApoE, IGF2, and IGFBP2) that increased transcription of synaptic genes in human neurons cultured on MEFs. Various factors were produced as human proteins in HEK293 cells (names reflect gene symbols; vectors and procedures are as described in previously in (37)) and added to human neurons on MEFs at D10 and incubated for 48 hours, before the harvest at D12 for qRT-PCR on indicated synaptic marker genes.

